# Orthogonal perturbation of sulfur availability reveals antibiotic-induced synthetic lethality in *Mycobacterium tuberculosis*

**DOI:** 10.64898/2026.07.28.741166

**Authors:** Vaibhav Kumar Nain, Vishawjeet Barik, Garhima Arora, Manitosh Pandey, Rahul Pal, Anjali Mishra, Ranjan Kumar Nanda, Samrat Chatterjee, Amit Kumar Pandey

**Affiliations:** Mycobacterial Pathogenesis Laboratory, Centre for Tuberculosis Research, BRIC-Translational Health Science and Technology Institute, Haryana, India; Complex Analysis Group, Computational and Mathematical Biology Centre, BRIC-Translational Health Science and Technology Institute, Haryana, India; Translational Health Group, International Centre for Genetic Engineering and Biotechnology, New Delhi, India

**Keywords:** sulfur acquisition, metabolic remodelling, CRISPRi, vulnerability and synthetic lethality

## Abstract

Despite its essential role in growth, the impact of sulfur limitation on mycobacterial physiology and antibiotic responsiveness remains poorly understood. Here, we combined chemical-genetic perturbations and infection models to investigate the consequences of sulfur limitation in *Mycobacterium tuberculosis (M. tuberculosis)*. We report that sulfur limitation restricted bacterial growth by rewiring metabolism, resembling nutrient starvation. CRISPR interference of the sulfate transporter (ST) revealed severe vulnerability and sensitised *M. tuberculosis* to anti-TB drugs. Inhibition of H_2_S production in macrophages further compromised the fitness of ST-deficient *M. tuberculosis*, indicating that the bacteria depend on host-derived sulfur during infection. Importantly, preventing sulfur acquisition attenuated both the drug-sensitive H_37_Rv and the drug-resistant clinical strain of *M. tuberculosis,* while enhancing the efficacy of Isoniazid (INH) in the murine model, resulting in sterilisation of infected lung tissues. Combined, our findings suggest that restricting sulfur uptake highlights a therapeutic vulnerability that enhances the efficacy of existing anti-TB drugs.

## Introduction

Despite decades of medical advancement, Tuberculosis remains the leading cause of mortality attributed to a single infectious agent, *Mycobacterium tuberculosis* (*M. tuberculosis*). The disease remains a global health emergency with approximately 10.7 million new cases and 1.23 million deaths reported globally in 2024. Consequently, *M tuberculosis* remains among the top bacterial priority pathogens on the WHO list (*1*, *2*). Alleviation of the above global health crisis warrants continuous discovery of novel drug scaffolds and development of innovative strategies to potentiate the efficacy of the existing anti-TB regimens.

To maintain its niche within the host microenvironment, *M. tuberculosis* carefully modulates its virulence, striking an evolutionary balance that prolongs infection to maximise transmission opportunities. This adaptation renders *M. tuberculosis* dependent on the host-derived macro- and micronutrients for both its survival and disease progression (*3*). Non-metallic elements such as nitrogen (N) (*4–8*) and phosphorus (P) (*9*, *10*) are important for *M. tuberculosis* during host infection, metabolism, and survival under host-induced stress (*11–13*). Likewise, sulfur is required for the biosynthesis of metabolites central to several metabolic pathways, critical enzymatic reactions, virulence factors, and redox homeostasis and energy metabolism (*14*, *15*).

*M. tuberculosis* can acquire sulfur mainly in the form of inorganic sulfate (or thiosulfates), organic forms such as S-containing amino acids (cysteine and methionine) or as hydrogen sulfide (H_2_S). Uptake of inorganic sulfate is aided by an ATP-binding cassette (ABC) transporter, CysTWA1-SubI **(Fig. S1)** (*14*, *16*, *17*), and is conserved across other bacterial species (*18*, *19*). In contrast, H_2_S, being gaseous, can readily traverse biological membranes passively and regenerate the bacterial thiol pool. In fact, host-generated H_2_S is required for the survival of *M. tuberculosis* during infection in mice (*16*, *17*). No dedicated transporters for cysteine and methionine have been identified in *M. tuberculosis*; however, a *Tn* insertion mutant of *rv3245c* homolog in *M. bovis* BCG was found to be a methionine auxotroph (*18*). Unlike bacteria, mammalian cells must acquire sulfur-containing metabolites, such as cysteine and methionine, from the surrounding medium. Cysteine is decomposed into pyruvate, ammonia and hydrogen sulfide, which is further oxidised to sulfate. As expected, during *M. tuberculosis* infection of a host macrophage, all these sulfur sources support its growth and survival **(Fig. S1)** (*19*).

The role of sulfur limitation in mycobacterial physiology, pathogenesis, and druggability has not been addressed systematically. This led us to hypothesise that targeting sulfur acquisition could impair the ability of *M. tuberculosis* to grow and infect host cells, and that co-treatment with existing anti-tubercular drugs could yield improved therapeutic outcomes. In this study, we have shown that sulfur deficiency leads to rewired bacterial metabolism and replication, characteristic of nutrient-starved *M. tuberculosis*. By targeting the sulfate transporter using CRISPRi, we demonstrated that *M. tuberculosis* was attenuated in growth and survival under both *ex vivo* and *in vivo* conditions. Finally, as previously suggested but not reported, abrogation of sulfur acquisition enhanced sensitivity to anti-TB drugs, improved the efficacy of Isoniazid in *M. tuberculosis* drug-sensitive strain, and reduced the burden of clinical INH^R^ isolate in murine lungs.

## Results

Sulfur limitation arrests the replication of *M. tuberculosis*

To investigate the consequence of sulfur limitation on the growth of *M. tuberculosis*, we prepared a sulfur-free (minuS) minimal medium and measured the total sulfur content to confirm sulfur depletion **(Fig. S2A)**. ^34^S (total sulfur) in the S-depleted medium was reduced significantly by ∼5 times compared to that of the medium containing sulfate salts **(Fig. S2B)**. We observed that the growth of *M. tuberculosis* and *M. smegmatis* was abrogated in a sulfur-free medium **(Fig. 1A and Fig. S2C),** indicating that sulfur availability is essential for mycobacterial growth *in vitro*. *M. tuberculosis* growing in a sulfur-lacking medium revealed ∼4-fold reduced total intracellular sulfur levels (^34^S) relative to those grown in the presence of sulfur, while no changes in other essential metal ions assayed, such as Magnesium (Mg), were observed **(Fig. 1A and Fig. S2D)**. To assay intrabacterial sulfate levels, we used a genetically encoded biosensor, Thyone, that exhibits a decrease in signal in the presence of sulfate ions (*20*). Thyone was fused to tagRFP for normalisation of the intensiometric response to sulfate ions and readout from the new ratiometric form, herein referred to as *mThyone* (Mycobacterial Thyone), was expressed as a GFP/RFP (Green/Red) ratio. *M. tuberculosis* expressing *mThyone* cultured in medium without sulfur exhibited a high Green/Red ratio in S-depleted bacteria compared to those grown in the presence of sulfur **(Fig. 1B)**. Supplementation of S-starved bacteria with different sources of sulfur (inorganic sulfate and amino acids) rescued bacterial growth, suggesting that *M. tuberculosis* is capable of salvaging sulfur from the amino acids - cysteine and methionine, in addition to inorganic sulfate **(Fig. S2E)**. Further, we sought to determine bacterial growth kinetics using the replication clock plasmid, pBP10 (*21*). *M. tuberculosis* harbouring the plasmid was cultured in media containing or lacking sulfur, and plasmid retention was calculated. The growth of *M. tuberculosis*, as expected in PluS medium, was rapid, with a high net growth and replication rate, leading to a rapid loss of the plasmid over time. Replication stalled in S-lacking bacteria, enabling them to retain the plasmid over time (∼70% of cells), with a stable rate of plasmid loss and no net growth or replication. Exogenous supplementation with ammonium sulfate (AS) increased replication and net growth rate, leading to plasmid loss **(Fig. S3, A to C)**. Interestingly, we observed that sulfur-starved bacteria with low or no replication rate showed a very low death rate as well, even in the late log-stationary phase (Day 12 onwards), indicating that the cells entered a non-replicating state. This phenotype was reversible as supplementing the medium with exogenous sulfate triggered a rapid resurgence in both replication and death rates. **(Fig. S3, D and E)**.

**Fig. 1:**
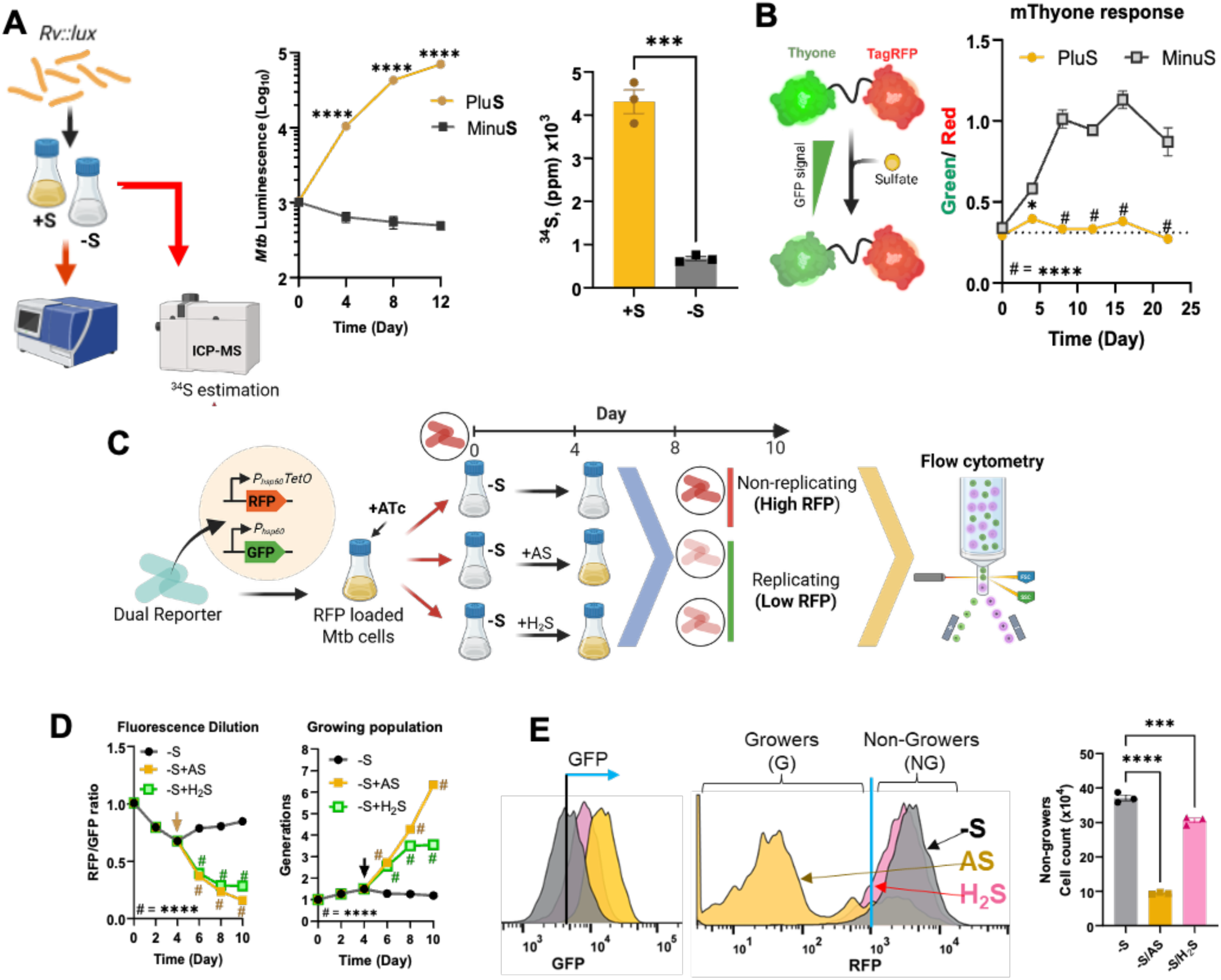
Sulfur-limitation curtails *in-vitro* replication of *M. tuberculosis*. **(A)** Growth kinetics of *M. tuberculosis* in chemically defined minimal medium, either lacking or containing inorganic sulfate(s) and ICP-MS based estimation of total sulfur content (^34^S) inside bacterial cells grown in respective media **(B)** Probing intrabacterial sulfate levels in *M. tuberculosis* using genetically encoded ratiometric sulfate biosensor mThyone. **(C)** Schematic representation of the fluorescence dilution assay for analysis of non-growing bacterial cells upon sulfur starvation by flow cytometry. Dilution of RFP indicates the growth in bacterial population. **(D)** Time-dependent kinetics of fluorescence dilution and actively growing cells in terms of generations. **(E)** Determination of grower and non-grower populations under S-starvation condition using fluorescence dilution as the replication marker. Data are the averages of the three data points indicated by dots and representative of at least two to three independent experiments. Error bars correspond to standard error of mean (SEM). Statistical significance was assessed by two-way ANOVA (Sidak’s multiple comparisons test) for (A) growth kinetic where ****P<0.0001; Unpaired Student’s t-test for ICP-MS where ***P = 0.0002; two-way ANOVA (Sidak’s multiple comparisons test) for (B) where *P = 0.01 and ****P<0.0001; ****P<0.0001 and two-way ANOVA (Dunnett’s multiple comparisons test) for (E) where ****P<0.0001 and one-way ANOVA (Dunnett’s multiple comparisons test) for (E) where ***P = 0.0006.

Since non-replication is associated with the emergence of a non-grower (NG) population of bacteria within the same culture, we sought to determine the percentage of non-growers in the presence or absence of sulfur. To do this, we employed single-cell fluorescence dilution (SCFD) analysis using a dual reporter system with constitutive GFP and *tet-*inducible RFP expression. Non-growing (NG) bacteria retain high RFP intensity, whereas it is diluted in the growing (G) population, resulting in low RFP intensities (*22*, *23*). *M. tuberculosis* harbouring dual reporter was ‘pre-loaded’ with RFP and cultured in sulfur-free medium. Subsequently, it was supplemented with either AS or H_2_S, and cells were analysed by flow cytometry **(Fig. 1C)**. For SCFD analysis, all GFP+ bacteria were gated, and RFP intensities were determined in cells grown in the absence of sulfur and those later supplemented with AS or H_2_S **(Fig. S4A)**. As expected, bacterial cells grown in the absence of sulfur retained high RFP until the addition of AS or H_2_S, when the dilution of RFP occurred as indicated by the right-to-left shifts in signal intensities **(Fig. S4B)**. FD over the course of time showed that supplementation of AS or H_2_S led to a rapid increase in generations as a consequence of rescued growth and replication leading to reduced RFP/GFP ratio, while both factors – generations and RFP/GFP remained unchanged during the assessment period in S-starved cells **(Fig. 1D).** Based on the RFP content, the percentages of growers and non-growers in S-depleted or supplemented (AS or H_2_S) cultures were determined. For instance, on day 6, S-starved cultures had more NG bacteria, which were reduced by ∼4-fold and ∼1.3-fold upon AS or H_2_S supplementation, respectively, with simultaneous increased GFP expression as an indicator of their physiological state. Growers, being physiologically and metabolically more active, expressed high GFP (indicated by left-to-right shift) as compared to the non-growers **(Fig. 1E)**. In sum, our findings indicate that the absence of sulfur arrests *M. tuberculosis* in a non-replicating state, which is reversible upon sulfur supplementation.

### Gene expression analysis of sulfur-starved *M. tuberculosis* revealed rewiring of crucial cellular processes

To gain an overview of how *M. tuberculosis* rewires its transcriptomic landscape and copes with S-starvation, we cultured the bacteria in the respective media and performed RNA sequencing followed by gene expression analysis **(Fig. 2A)**. We observed that *M. tuberculosis* exhibited a distinct expression profile, with several genes downregulated under S-starvation **(Fig. 2B)**. Majorly, the expression of genes involved in intermediary metabolism and respiration, followed by conserved hypothetical genes and cell wall-associated genes/proteins, was altered under S-starved conditions compared to a standard medium containing sulfur **(Fig. 2C)**. Change in gene expression upon S-starvation was associated with GO terms and KEGG pathways that were significantly affected - redox homeostasis, electron transport chain, starvation, protein folding, translation, ribosome assembly, etc. **(Fig. 2D and Fig. S5, A and B)**. Upon closer inspection, we observed that genes encoding Type I NADH dehydrogenase (NDH-1), ATP synthase, Cytochrome *bc1* reductase, Cytochrome *c* reductase, Cytochrome *bd* oxidase, Succinate dehydrogenase, ferredoxins, and other components of the bacterial electron transport chain were significantly downregulated **(Fig. 2E and Fig. S6A)**. Next, genes encoding ribosomal protein subunits – L, M and S showed downregulated expression, as did chaperones involved in protein folding and the protein degradation system **(Fig. 2E and Fig. S6B)**. Genes encoding sulfate transporter assembly, cysteine and methionine metabolism were upregulated **(Fig. 2E)**.

**Fig. 2:**
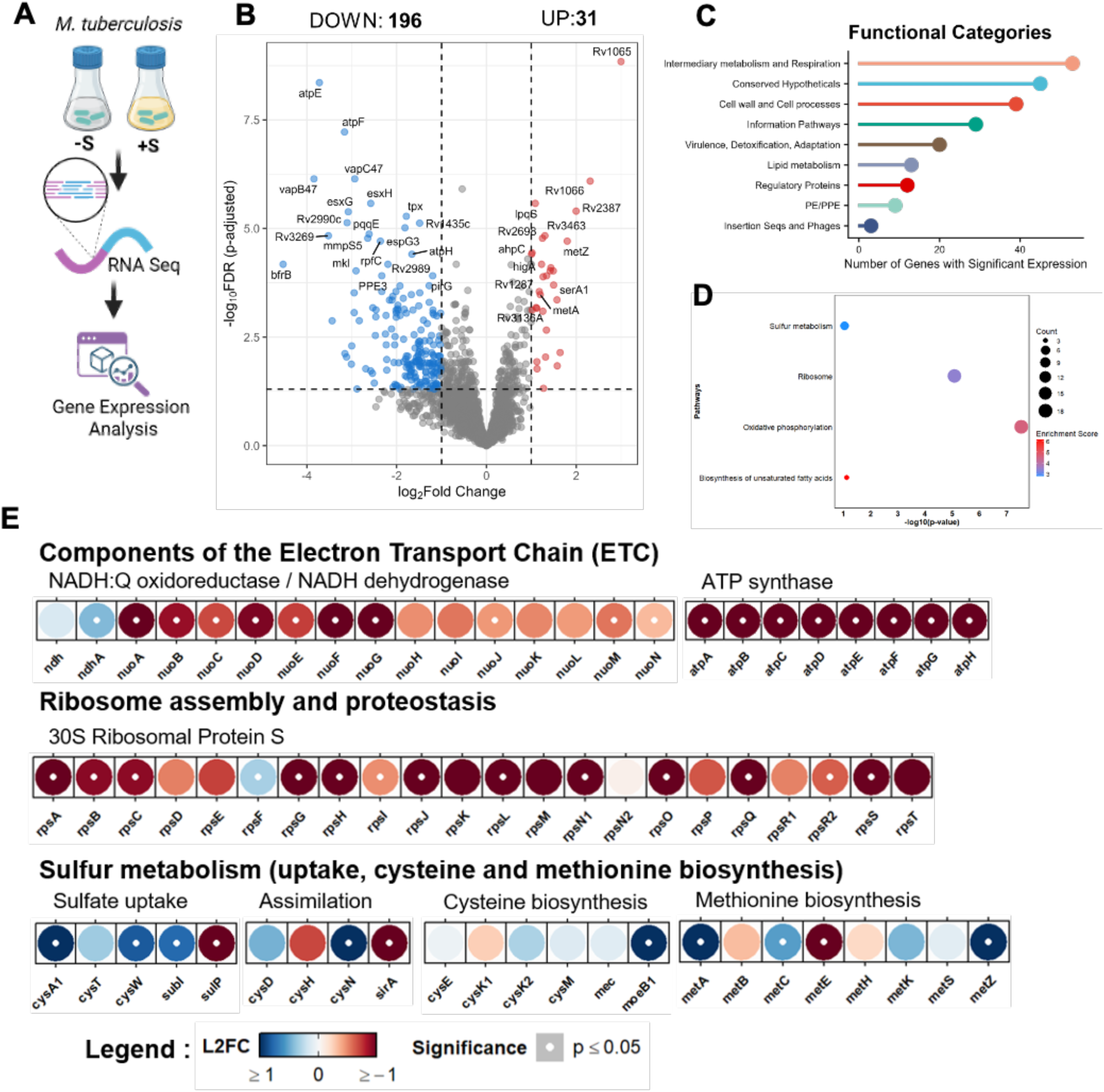
Gene Expression analysis of sulfur-starved *M. tuberculosis*. **(A)** Graphical representation of the transcriptomics experimental plan. *M. tuberculosis* was grown in minimal medium (MM), either supplemented or not with sulfate-containing salts. RNA sequencing was conducted to identify the differentially expressed genes (DEGs) under S-free vs. S-rich conditions. **(B)** Volcano plot depicting gene expression profile of *M. tuberculosis* under S-free vs S-rich conditions. DEGs were identified based on expression (log_2_ Fold change) cut-off value set at ±1 (dotted vertical line) and FDR-adjusted *p*-value ≤0.05 (dotted horizontal line). **(C)** Functional categories of the genes with significantly altered expression. **(D)** Pathway analysis of the genes with altered expression. **(E)** Feature expression heatmaps of key functional pathways. The colour of each circle represents the expression level of genes as log_2_ fold change (L2FC), where |L2FC|≥1 was set cut-off for change in gene expression, denoting up or down-regulation, and the white dot represents *p*-value (Significance) ≤0.05. Pathways depicted are - Components of the Electron Transport Chain (ETC) – NADH dehydrogenase, ATP synthase; Ribosome assembly and protein homeostasis and Sulfur metabolism – sulfur acquisition and assimilation.

Additionally, genes involved in redox homeostasis (transcription factors - *whiB3, whiB6, whiB7,* peroxidases, thioredoxins and mycothiol were upregulated in response to increasing cytosolic oxidative stress **(Fig. S6F)**. Lastly, genes critical for cell division (*ftsZ*, *wag31*, *wag32*, *pirG*, etc.) were downregulated, potentially resulting in reduced cell division under S-limitation conditions. Other physiological and/or metabolic processes affected include glycolysis (*fba*, *gap* and *pgk*), TCA cycle (*aceE, acn*, *citA*, *dlaT* and *pckA*), glutamate biosynthesis (*gltA2*, *gltB* and *gltD*), transcription (*rpoA*, *rpoZ*, *nusG,* and *rho*), and fatty acid metabolism (acetyl-CoA acetyltransferases), which were down-regulated in concordance with our *in vitro* findings **(Fig. S6, C and G to J)**. In sum, sulfur deprivation influences bacterial energy metabolism, proteostasis and other essential physiological processes while also signaling the need for increased sulfur acquisition and regeneration of the total thiol pool within cells in response to a drop in intrabacterial sulfur levels.

### Sulfur limitation perturbs core metabolic pathways in *M. tuberculosis*

Transcriptomic profiling of S-starved *M. tuberculosis*, encompassing expression data for 4008 unique genes, revealed that approximately 25% of the DEGs were involved in metabolic processes. Therefore, to gain insight into the metabolic reprogramming that bacteria undergo upon sulfur deprivation, we overlapped the preprocessed gene list with the 1,011 unique metabolic genes in the *iEK1011* genome-scale metabolic model (GSMM) (*24*, *25*). We observed that expression data for 1,000 metabolic genes across both conditions covered ∼90% of the available reactions. We integrated the expression data into the *iEK1011* metabolic model and converted gene expression values to reaction-level activity scores using the model’s gene-protein-reaction (GPR) rules. Furthermore, context-specific metabolic models (S-deprived and S-rich) were prepared using the E-Flux method (*26*), which uses reaction activity scores to constrain the upper and lower bounds of reaction fluxes **(Fig. 3A)**. Next, to explore the infinite steady-state solution space without optimisation, we used the GP sampler (*27*, *28*), which generates feasible steady-state flux vectors that satisfy all network constraints. We sampled the flux space and identified the mean flux vector as the representative solution. Convergence analysis showed that the mean flux vector stabilised at sample sizes of 12,380 for the control and 30,950 for the test condition. **(Fig. 3B)**.

**Fig. 3:**
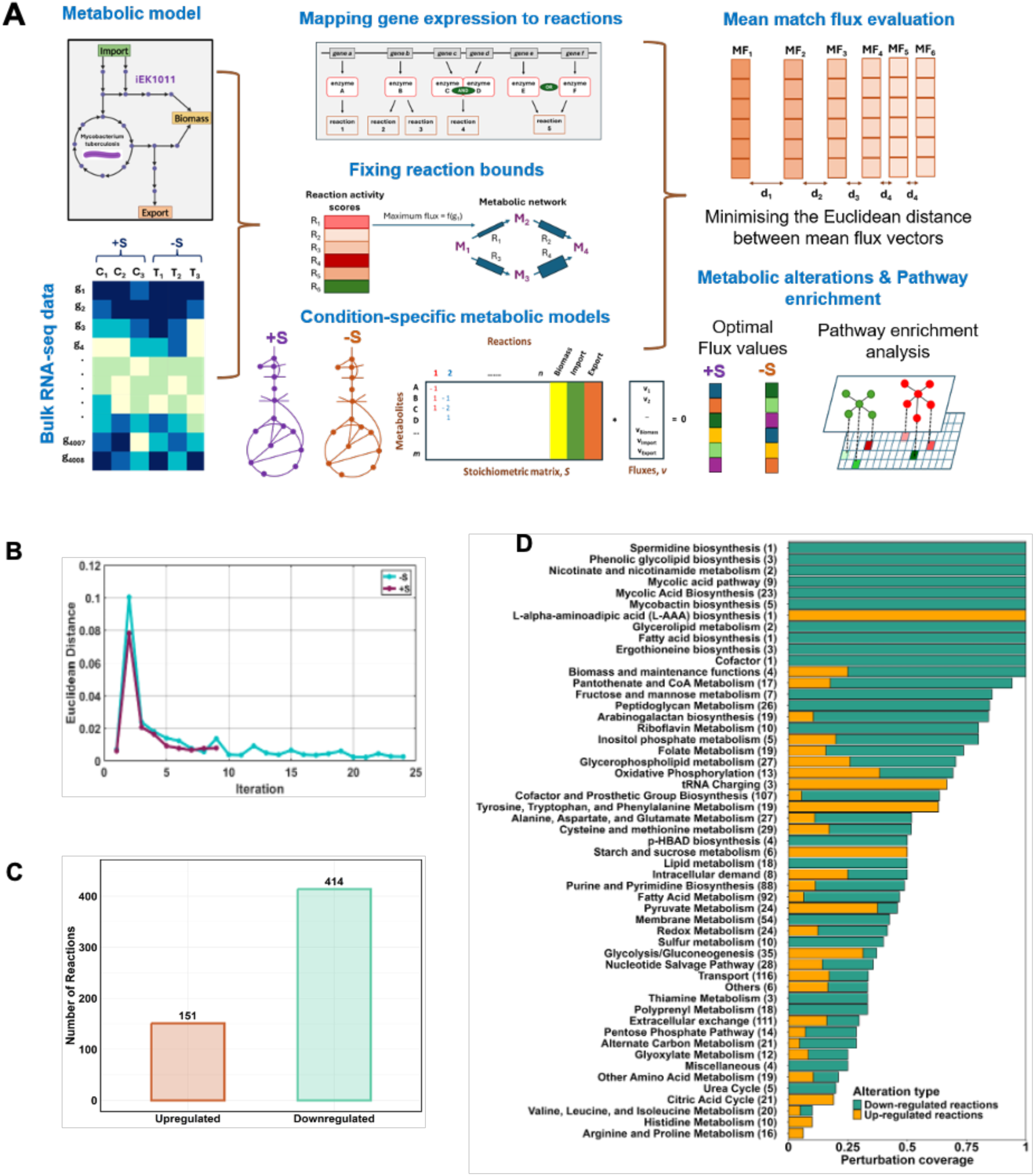
Integrated analysis of differential gene expression, flux convergence, and pathway perturbations under sulfur limitation. **(A)** Graphical representation of the workflow involved in the genome-scale metabolic modelling. **(B)** Convergence analysis of the flux sampling using the GP sampler for Sulfur-rich and sulfur-limited conditions. The Euclidean distance between mean flux vectors from successive iterations decreases and converges to a representative steady-state flux distribution. **(C)** The bar plot represents the number of perturbed reactions. **(D)** Pathway perturbation analysis based on perturbed reactions. Bars represent perturbation coverage for each metabolic pathway, with green indicating the proportion of downregulated reactions and orange indicating the proportion of upregulated reactions. The total number of reactions in each pathway is indicated in parentheses.

We next identified reactions perturbed in the test (sulfur-deprived) versus control (sulfur-rich) conditions, using a 1.5-fold cutoff on flux fold-change values for reactions common to both condition-specific models. Reactions that met this criterion but showed opposite flux directions between the two conditions were removed from further analysis. Using this approach, we identified 151 upregulated and 414 downregulated reactions in the test condition relative to the control (**Fig. 3C**). Pathway analysis of the differentially regulated reactions between test (sulfur-deprived) and control (sulfur-supplemented) conditions in *M. tuberculosis* was performed using annotations from the iEK1011 metabolic model database.

High-coverage pathways associated with cell wall lipid biosynthesis (mycolic acid, phenolic glycolipid, and peptidoglycan metabolism); membrane metabolism; and polyprenyl metabolism were found to be altered and downregulated (**Fig. 3D**). These pathways are essential for the bacterial cell wall structure and virulence, and their reduced activity in sulfur-deficient environments suggests either delayed growth or a stress-adapted state. Significant disruption of central carbon metabolism-related pathways, including glycolysis/gluconeogenesis, the citric acid cycle, oxidative phosphorylation, and the pentose phosphate pathway, was also identified, with metabolic rewiring characterised by a mixture of upregulated and downregulated reactions under sulfur-deprived conditions. Sulfur utilisation pathways, including cysteine and methionine metabolism, as well as redox processes, were also significantly perturbed, suggesting that the bacterium is struggling to maintain redox balance. Some reactions were significantly upregulated, including tRNA charging, cofactor and prosthetic group biosynthesis, nucleotide salvage pathways, intracellular demand or transport and parts of amino acid metabolism, indicating activation of compensatory maintenance mechanisms and adaptation to stress conditions. Overall, in line with our earlier findings, the pathway enrichment analysis showed that in the absence of sulfur, the bacterium is under nutritional stress, as indicated by the alteration of reaction rates, possibly leading to changes in levels of metabolites critical for growth and virulence, and, simultaneously, the pathogen appears to be upregulating pathways involved in the mitigation of the above stress.

### Sulfur limitation affects respiration and stalls biosynthesis in *M. tuberculosis*

Metabolic flux predictions and transcriptome analysis suggested that the lack of sulfur rendered *M. tuberculosis* non-replicating by influencing metabolic processes, including respiration, biomass production, and transcriptional/translational activity. To validate this experimentally, we cultured *M. tuberculosis* in S-rich and S-free media and subsequently determined total protein and ATP abundances. As expected, compared with cells grown in sulfur, S-starved bacteria showed ∼5-fold and ∼3-fold lower total protein and ATP content, respectively **(Fig. 4A)**. Next, using the ratiometric biosensor Peredox-mCherry (*29*), we determined the metabolic state of S-starved *M. tuberculosis* and found it to be stalled, as indicated by a higher NADH/NAD+ ratio, suggesting non-recycling of NADH to NAD+ and vice versa **(Fig. 4B)**. Further, we determined the transcriptional/ translational activity in *M. tuberculosis* upon S-starvation. To do this, we again used the dual-reporter system, with constitutive GFP expression as a marker of bacterial growth and Tet-inducible RFP as a marker of transcriptional/translational activity. *M. tuberculosis* dual reporter strain was grown in media ±S and supplemented with Anhydrotetracycline chloride (ATc) to induce expression of RFP. We observed lower and sustained RFP expression (transcriptional-translational activity) in S-starved *M. tuberculosis*. Supplementation of AS to S-starved culture at day 8 led to an increase in both bacterial growth (GFP) and transcriptional activity (RFP) **(Fig. 4C)**. Reduced biomass and metabolic activity indicate non-utilisation of available nutrients for biosynthesis. In line with this, we performed a meta-analysis in which we correlated the transcriptome of carbon (or nutrient)-starved (*30*) with that of S-starved *M. tuberculosis* **(Fig. S7A)**. Interestingly, among the genes commonly downregulated under both conditions, most were primarily involved in respiration, information, cell wall, redox, and stress homeostasis, indicating that S-limitation might induce a reversible starvation-like state despite nutrient availability **(Fig. S7B)**. Likewise, we observed that the attenuation of growth in the absence of sulfur was independent of the carbon source used in the medium. *M. tuberculosis* grown in S-free medium with different carbon sources – glycerol, cholesterol, or propionate showed no growth until supplementation with a sulfur source, indicating that sulfur is essential for the utilisation of carbon sources **(Fig. S7C)**. These findings emphasise that sulfur limitation impedes growth by stifling the metabolic and physiological activity of *M. tuberculosis,* thereby causing the bacterium to enter a metabolically quiescent, non-replicating state.

**Fig. 4:**
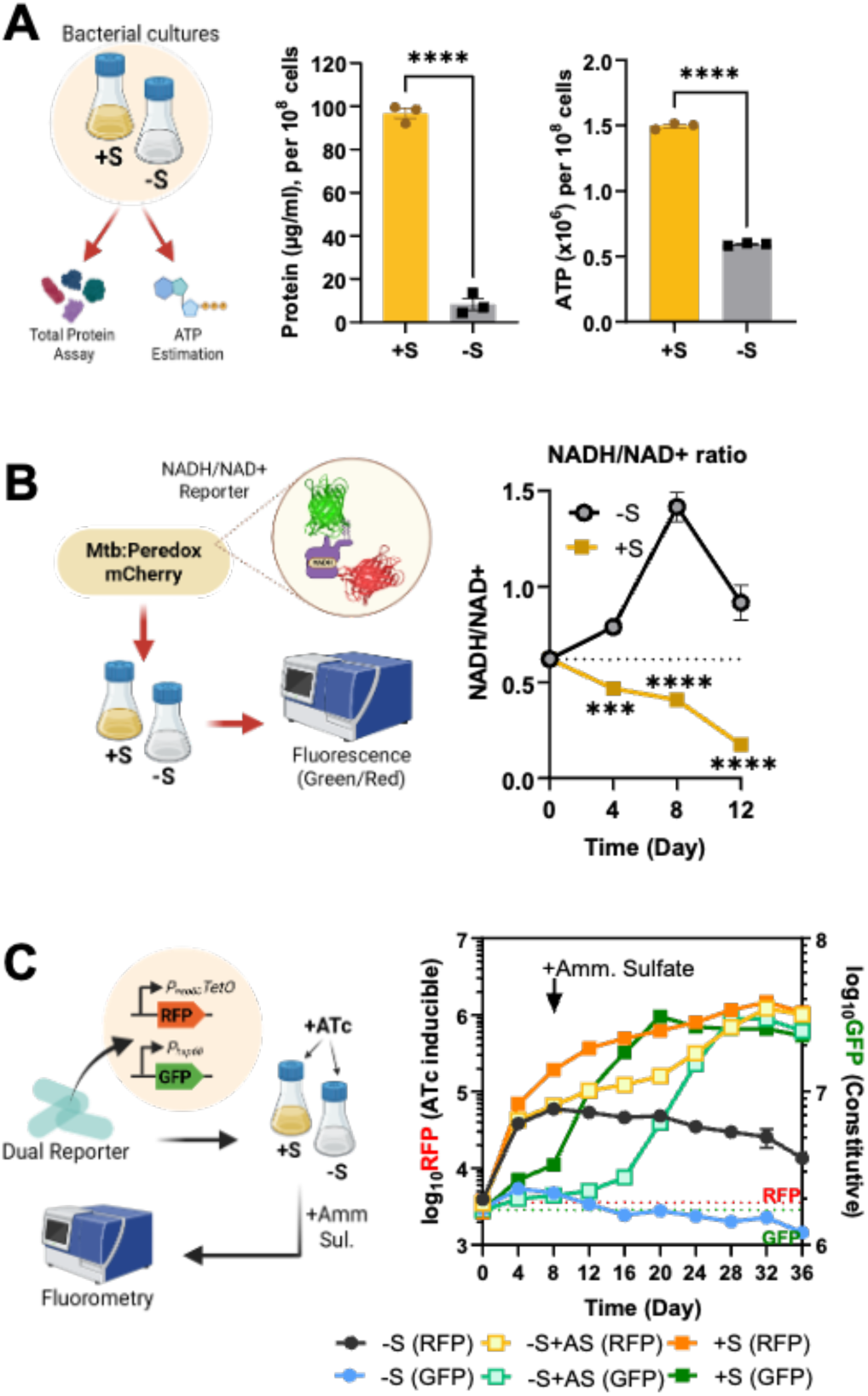
S-limitation leads to metabolic rewiring in *M. tuberculosis*. **(A)** *M. tuberculosis* grown in PluS and MinuS media was assayed for total protein and ATP levels (using Bactitre Glow). **(B)** Estimation of NADH/NAD+ levels in sulfur-starved *M. tuberculosis* using ratiometric biosensor, Peredox-mCherry. **(C)** Transcriptional-Translational Activity (TTA) assay to determine the physiological activity of sulfur-starved *M. tuberculosis* using a dual reporter strain in terms of active RNA and protein biosynthesis. Data are the averages of the three data points indicated by dots and representative of at least two to three independent experiments. Error bars correspond to standard error of mean (SEM). Statistical significance was assessed by Unpaired Student’s t-test for (A), where ****P<0.0001; two-way ANOVA (Sidak’s multiple comparisons test) for (B), where **P = 0.00024; ***P = 0.0008 and ****P<0.0001.

### Genetic depletion of sulfur acquisition reveals vulnerability in *M. tuberculosis*

S-starved *M. tuberculosis* exhibited increased expression of sulfate-acquisition genes, prompting us to generate a genetic model with restricted inorganic sulfur uptake to facilitate further investigation of the role of sulfate acquisition in bacterial pathogenesis. The sulfate transporter in *M. tuberculosis* is encoded by the operon *cysTWA1-subI* (*rv2400c-rv2387c*), an ABC transporter (*31*) and *sulP* (*rv1739c*), a non-specific solute transporter (*32*) **(Fig. 5A)**. Further to this, since our goal was to broadly abolish sulfate import by the bacterium, a combined rather than individual genes/ transporters were targeted. To achieve this, *cysT*, *cysW*, and *sulP* were knocked down in tandem using CRISPRi (*33*, *34*) **(Fig. 5B)**, and target genes’ expression was verified by RT-qPCR to confirm the knockdown **(Fig. S8A)**. A sulfate transporter(s) knockdown mutant of *M. tuberculosis* (hereafter referred to as ST_KD_) was resistant to selenium and chromate in concordance with previous findings **(Fig. S8B)** (*31*, *35*). ATc titration revealed high *in vitro* vulnerability of sulfate uptake in *M. tuberculosis,* wherein low-level suppression of the transporter (ATc IC_50_ ∼10 ng/ml) had severe consequences for bacterial growth **(Fig. S8C)**. The growth rate of *M. tuberculosis* is not affected by ATc concentrations as high as 1-5 µg/ml, confirming that the observed vulnerability phenotype reflects target gene(s) knockdown rather than the direct effect of ATc (*36*). Likewise, ST_KD_ failed to grow in a standard culture medium containing multiple sulfate sources **(Fig. 5C and Fig. S8D),** similar to *M. tuberculosis* grown in S-free medium. Intracellular sulfate levels in ST_KD_, when probed with *mThyone*, were lower, as indicated by a high Green/Red ratio **(Fig. 5D)**. The growth defect in ST_KD_ was rescued by methionine, cysteine and NaHS **(Fig. S8E)**. Additionally, inhibition of sulfate uptake led to reductions in ATP (∼2-fold) and total thiol content (∼2.5-fold), with a comparable membrane permeability **(Fig. S8F)**. Further, ST_KD_ harbouring Peredox-mCherry showed a rise in NADH/NAD+ ratio in response to ATc-dependent suppression of the transporter **(Fig. S8G)**. The proteome analysis of *M. tuberculosis* with and without a sulfate transporter highlighted several differentially expressed proteins that majorly affect sulfur metabolism, respiration, redox homeostasis, purine and amino acid metabolism, and ribosome biosynthesis, like that of S-starved *M. tuberculosis* **(Fig. 5E and Fig. S9A)**. Given a similar signature, upon comparison with the transcriptomic profile of S-starved *M. tuberculosis*, we identified 43 genes common to the core physiology of the bacterium – respiration, ribosome, redox, etc. **(Fig. S9B)**. In sum, a dose-dependent CRISPRi demonstrated functional vulnerability to sulfur depletion in *M. tuberculosis,* resulting in slow replication and growth rates, along with low ATP and thiol content, the same physiological hallmarks and shortcomings as those of sulfur starvation in *M. tuberculosis*.

**Fig. 5:**
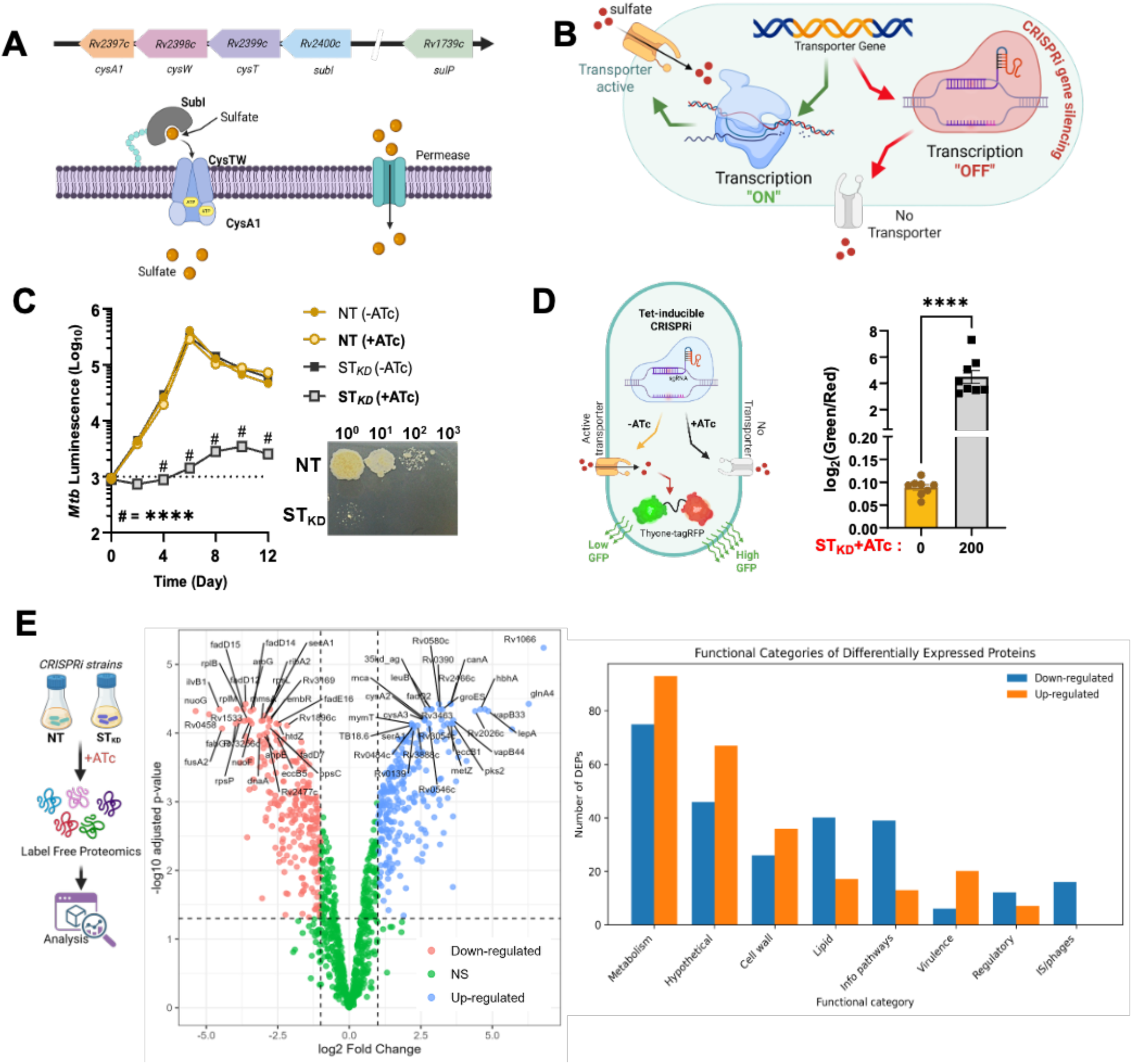
Inhibition of sulfate transporter(s) leads to attenuation in *M. tuberculosis.* **(A)** Graphical representation of the genetic locus *(rv2397c-rv2400c)* encoding the ABC transporter for sulfate uptake – *cysTWA1-subI* and permease, *sulP* encoded by *rv1739c*. **(B)** Schematic representation of the transcriptional knockdown of sulfate transporters by CRISPR interference. **(C)** Growth kinetics of sulfate transporter(s) knocked down (ST_KD_) *M. tuberculosis,* in contrast with the Non-targeting (NT) strain in sulfur-rich regular medium. **(D)** In-vitro sulfate levels assay using mThyone in NT and ST_KD_ strains. **(E)** Proteomic analysis of Sulfate transporter knockdown (ST_KD_) strain and comparison of sulfur-starved *M. tuberculosis* transcriptomics and ST_KD_ *M. tuberculosis* proteomics. The data are averages of the three data points indicated by dots and are representative of at least two independent experiments. Error bars correspond to the standard error of the mean (SEM). Statistical significance was assessed by two-way ANOVA (Sidak’s multiple comparisons test) for (C), where ****P<0.0001 and unpaired Student’s t-test (Welch’s correction) for (D), where ****P<0.0001.

### Host-derived sulfide compensates for the sulfur limitation in *M. tuberculosis* **during infection**

To assess whether the inability to acquire inorganic sulfate compromises the virulence of *M. tuberculosis*, we used a macrophage infection model. ST_KD_ was either pre-depleted or not for knockdown of sulfate transport(s) prior to macrophage infection. The ability to infect macrophages was determined by intracellular CFU post infection, which was significantly reduced (∼1 log_10_) in the ST_KD_ pre-depleted strain compared to the NT (non-targeting) control strain, while knocking down the transporter only for the period of phagocytosis (i.e., 4 hours) resulted in (∼0.5 log_10_), revealing that acquisition of sulfate is required for the virulence of *M. tuberculosis* **(Fig. 6A)**. To determine the *in-vivo* pathogenesis of *M. tuberculosis* lacking a sulfate transporter, we infected mice with the ST_KD_ and induced transporter knockdown by administering doxycycline immediately post-aerosolisation and again 1-week post-infection, followed by CFU plating **(Fig. 6B)**. We observed a significantly reduced bacterial burden in bacteria lacking the transporter, with a more pronounced attenuation when sulfate acquisition is inhibited immediately post-infection, indicating that sulfate uptake is essential for pathogenesis and infection establishment **(Fig. 6C)**. Consequently, gross pathology of representative lungs from -Dox (wild-type) and +Dox (0-week knockdown) at week 3 post-infection showed improved lung pathology, with no visible granulomatous lesions, compared to wild-type *M. tuberculosis,* and was concordant with significantly reduced granuloma scores **(Fig 6D)**. We also observed a reduced bacterial burden in the spleen, with similar levels of attenuation in both groups (0-week and 1-week knockdown) **(Fig. 6E)**. Despite attenuation, *in vivo* growth of the ST_KD_ increased steadily, possibly due to host-generated H_2_S, which is readily available to bacteria upon infection (*16*, *17*, *37*). To test this, we infected THP1-derived macrophages with mycobacterial strains (NT and ST_KD_), treated them with the CSE (cystathionine-γ-synthase) inhibitor PAG (D-L-Propargylglycine), and monitored bacterial viability by CFU counts or bioluminescence. We find that, in concordance with previous reports, inhibiting H_2_S biosynthesis curtailed the growth of wild-type *M. tuberculosis*, whereas the ST_KD_ exhibited severe attenuation with a reduction in bacterial burden by ∼50% **(Fig. 6F)**. These findings confirm that *M. tuberculosis* requires both inorganic sulfate and host-generated H_2_S for infection and disease progression, and that co-inhibition of both effectively restricts bacterial growth in macrophages.

**Fig. 6:**
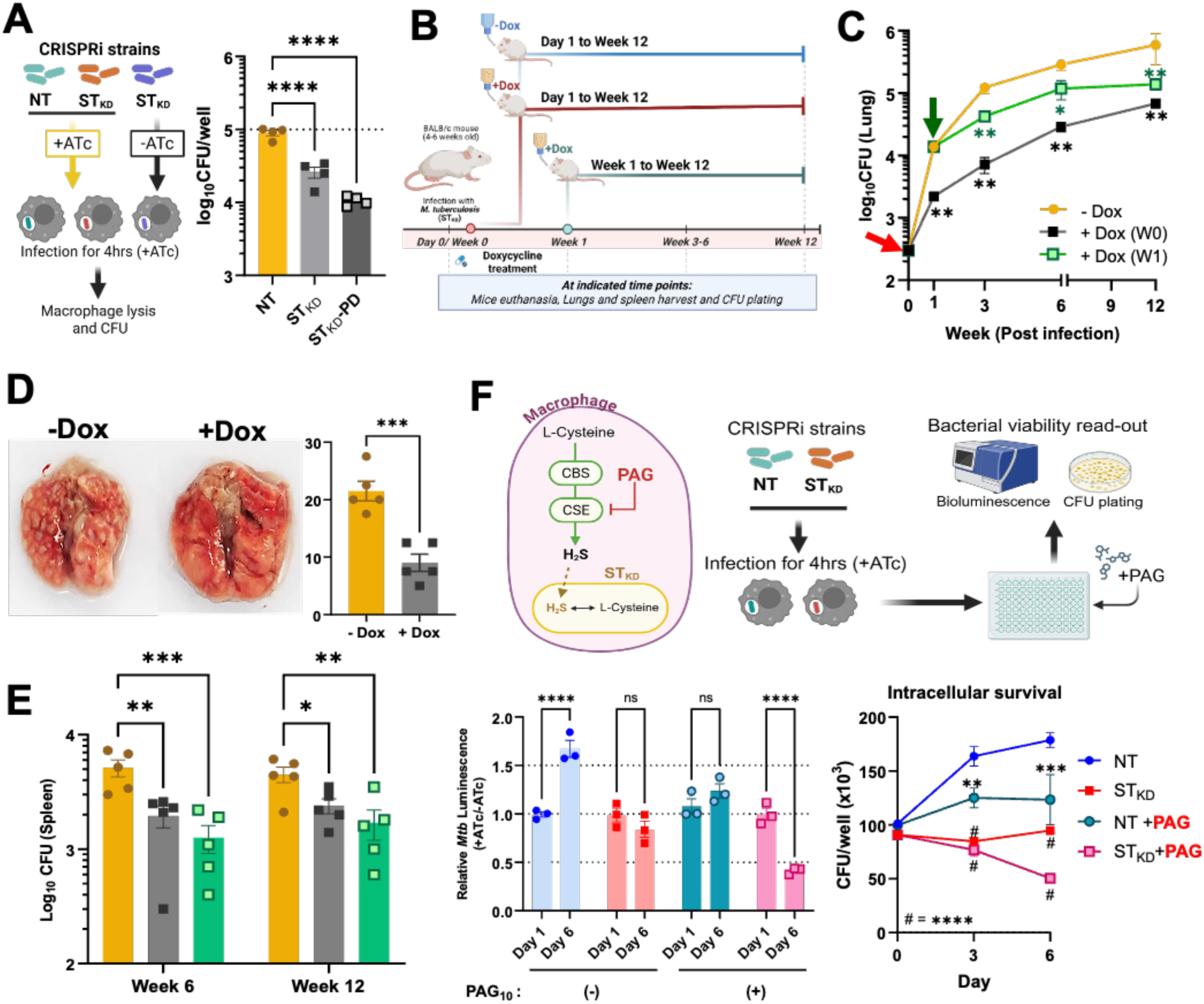
Acquisition of inorganic sulfur and host-derived H_2_S is essential for in vivo survival of *M. tuberculosis*. **(A)** Determining *ex-vivo* virulence of *M. tuberculosis* deficient in sulfate acquisition. ST_KD_ pre-treated or not with ATc were allowed to infect THP-1 macrophages compared to NT strain. **(B)** Diagrammatic representation of the experimental set-up to determine *in-vivo* pathogenesis of *M. tuberculosis* lacking sulfate acquisition machinery. Transcriptional knockdown of the sulfate transporter was initiated by doxycycline administration (2mg/ml) and bacterial burden was determined at indicated time-points. **(C)** Bacterial burden in mice lungs at indicated time-points. Red and green arrowheads indicate the start of doxycycline treatment at day 1 and day 7 post-infection respectively. **(D)** Representative images of gross pathology of lungs infected with ST_KD_ *M. tuberculosis,* either treated or not treated with doxycycline (inducer of CRISPRi knockdown). **(E)** Bacterial burden in infected spleens at indicated time-points. **(F)** H_2_S by macrophages acts as a sulfur source for *M. tuberculosis*. *Ex-vivo* infection model showing survival of wild-type and sulfur-starved *M. tuberculosis* inside macrophages upon treatment of PAG (Inhibitor of H_2_S biosynthesis) in the host. The data are averages of the three data points indicated by dots and are representative of at least two independent experiments. Error bars correspond to the standard error of the mean (SEM). Statistical significance was assessed by one-way ANOVA (Dunnett’s multiple comparisons test) for (A), where ****P<0.0001; and Mann-Whitney test for (C), where *P = 0.0317; **P = 0.0079; Unpaired Student’s t-test for (D), where ***P = 0.0006; two-way ANOVA (Dunnett’s multiple comparisons test) for (E), where *P = 0.0271; **P = 0.0003; ***P = 0.0001; two-way ANOVA (Sidak’s multiple comparisons test) for (F), luminescence based read out where, ****P<0.0001 and Dunnett’s multiple comparisons test for CFU based read out where, **P = 0.0085; ***P = 0.003 and ****P<0.0001

### Sulfur limitation imposes drug-induced synthetic lethality in *M. tuberculosis*

To investigate whether targeting sulfate uptake alters *M. tuberculosis* sensitivity to anti-tubercular drugs, the minimum inhibitory concentration (MIC) was determined after transporter knockdown. We observed that *M. tuberculosis* lacking the sulfate transporter (ST_KD_) is rendered sensitive to RIF, BDQ, EMB, and CFZ, with reduced IC_50_ values compared to those of wild-type (NT) bacteria, but not INH **(Fig. 7A)**. Next, to determine whether targeting sulfate acquisition has a cidal effect, NT and ST_KD_ were treated with different concentrations of ATc (100, 500, 1000ng/ml). We observed a significant reduction in fold survival of transporter-deficient *M. tuberculosis* at increasing ATc doses alone, compared with NT (wild-type) bacteria, which underlies the importance of sulfate acquisition in bacterial survival **(Fig. 7B and Fig S10A)**. Further, to assess whether inhibition of sulfate acquisition can potentiate the killing activity of anti-TB drugs, both NT and ST_KD_ were co-treated with ATc (100 ng/ml) and higher concentrations of drugs: RIF (10X), INH (10X), and BDQ (5X). Indeed, killing was potentiated, reducing ST_KD_ survival several-fold for all three drugs. We recorded a ∼1.5-2.5 log_10_ reduction in bacterial survival upon combinatorial treatment of ATc with RIF and BDQ, followed by INH **(Fig. S10B)**. Finally, we conducted an *in* vivo drug-efficacy study using the ST_KD_ strain of *M. tuberculosis* to assess the impact of inhibition of sulfate acquisition on antibiotic therapy. The standard dose of Isoniazid (INH; 10mg/kg body weight) was administered in combination with doxycycline (to knock down the sulfate transporter), and bacterial viability was monitored. We find that the group of mice infected with *M. tuberculosis* lacking transporter (+Dox) showed faster killing kinetics than the group that was still able to uptake sulfate from the host tissue (- Dox) and was eventually eliminated from the lungs by the end of 2 weeks of INH treatment, leading to a comparatively better therapeutic outcome **(Fig. 7C)**. In sum, the thiol pool in bacterial cells protects them from the cidal effects of antibiotics, and the absence of inorganic sulfur acquisition leads to efficient sterilization of bacteria within the tissue when treated in combination with INH.

**Fig. 7:**
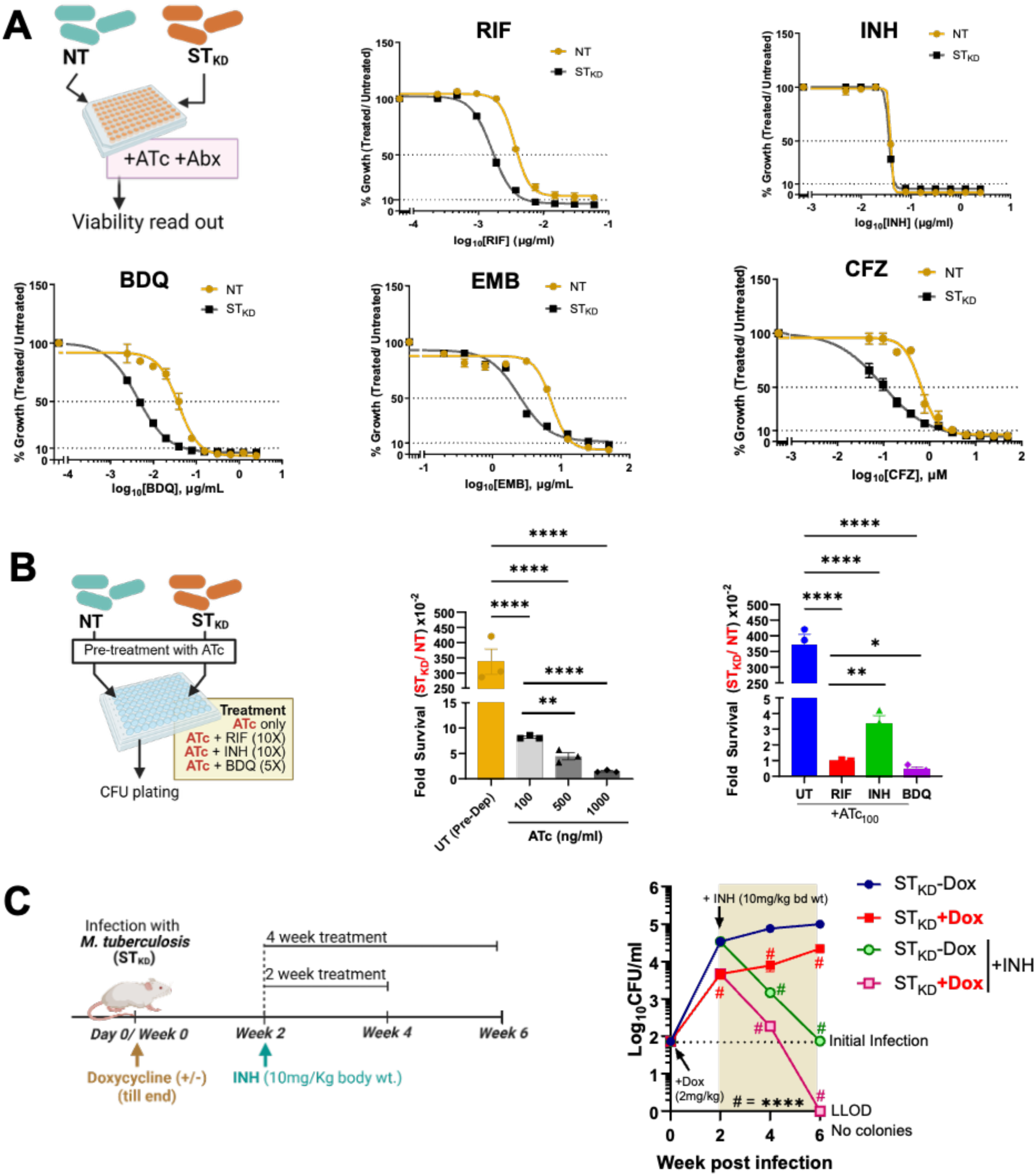
Sulfur-starved *M. tuberculosis* is sensitised to anti-TB drugs. **(A)** Schematic representation of chemical genetic interaction with anti-TB drugs – RIF, INH, BDQ, EMB and CFZ. The anti-tubercular activity is depicted as dose-response curves against wild-type (ATc 0) and sulfate transporter knockdown (ST_KD_) *M. tuberculosis* (ATc 100) **(B)** *In-vitro* drug susceptibility of *M. tuberculosis* lacking sulfate transporter to anti-TB drugs. **(C)** Experimental setup for determining the efficacy of anti-TB drug INH against sulfate-depleted *M. tuberculosis*. ST_KD_ *M. tuberculosis-*infected mice were either treated or not treated with doxycycline to induce knockdown of sulfate transporter(s). At the indicated time point, INH treatment was initiated, and bacterial clearance was assessed by determining bacterial CFU in lung homogenates from infected/treated animals. The data are averages of the three data points indicated by dots and are representative of at least two independent experiments. Error bars correspond to the standard error of the mean (SEM). Statistical significance was assessed by Ordinary one-way ANOVA for (B: ATc survival), where **P = 0.0016 and ****P<0.0001; (B: drug survival), where *P = 0.00371; **P = 0.0084 and ****P<0.0001 and two-way ANOVA (Dunnett’s multiple comparisons test) for (C), where ****P<0.0001.

### Drug-resistant *M. tuberculosis* demonstrates enhanced susceptibility to sulfur restriction

Clinical *M. tuberculosis* isolates acquire mutations that confer drug resistance at the expense of fitness (*38*), while non-resistance mutations in crucial metabolic networks can also influence bacterial survival and virulence (*39*). To investigate whether sulfur metabolism is compromised in clinical isolates of *M. tuberculosis*, we intended to identify mutations in genes involved in sulfur metabolism using the CRyPTIC open-source clinical data compendium (*40*, *41*). Sulfur uptake, processing, and assimilation are coordinated via multi-step metabolic reactions carried out by specific enzymes **(Fig. 8A)**. Our analysis revealed that mutations in sulfur metabolism genes were near-universal across the dataset, which is expected given the size of the gene panel; mutation frequencies nonetheless varied substantially by gene, with highest frequency observed in genes related to sulfate uptake, cysteine, and mycothiol biosynthesis. **(Fig. 8B)**. Further, we assessed the association of mutations in these genes with drug resistance or susceptibility. Interestingly, despite the fact that mutations in all genes existed in drug-resistant as well as sensitive isolates, those in *subI*, *cysW*, *cysK2*, *sahH* and *mshA* co-occurred at elevated frequency with multidrug and extensively drug-resistant profiles **(Fig. 8C).** It is important to note this pattern may partly reflect lineage-specific associated mutation accumulation rather than a resistance-specific mechanism. A few genes (*cysK2*, *cysK1*, *metH*, *cysW*, *mshA*, *subI*, *egtED*, etc.) showed a high frequency of non-synonymous mutations across the clinical isolates **(Fig. S11A)**. Non-synonymous mutations are mostly responsible for structural and functional changes at the protein level, possibly leading to *loss-of-function*. We looked at the identity of the most frequently occurring non-synonymous mutations in the panel of selected sulfur metabolism genes and found that *cysW* (G141A), *subI* (A76T), *mshA* (A187V and/or N111S), *cysK2* (A101S and/or G93S), *metH* (R539H), etc., were amongst the mutations occurring in drug-resistant profiles **(Fig S11B)**. Given the broad presence of non-synonymous mutations in sulfur metabolism genes across clinical isolates, including multidrug-resistant strains, we tested whether such isolates remain vulnerable to sulfate limitation directly. For this, we used a clinical INH^R^ isolate (XTB13-198, BEI Resources) (*42*) and ascertained INH resistance in the clinical strain relative to the INH-sensitive H_37_Rv. WGS analysis of XTB13-198 revealed mutations in *cysW*, *cysT, cysD*, *cysK2* and *metK* genes suggesting compromised sulfur uptake and assimilation into cysteine and methionine **(Fig. 8D)**. Next, we knocked down the sulfate transporter(s) in the clinical isolate using CRISPRi and observed in vitro growth attenuation, as confirmed by growth kinetics **(Fig. 8E)**. ATc titration in ST_KD_ strains of H_37_Rv and INH^R^ *M. tuberculosis* revealed that the clinical strain was more susceptible to sulfur depletion (∼100 times) than the laboratory strain **(Fig. 8F)**. Further, INH^R^::ST_KD_ was used to infect mice and bacterial clearance upon doxycycline administration showed a significant reduction (>2 log_10_) in bacterial burden in the knockdown group with no replication in host tissues **(Fig. 8G)**. These results indicate that drug-sensitive and resistant strains share vulnerability to sulfur limitation; hence, inhibiting sulfur acquisition can serve as a therapeutic strategy to effectively control drug-resistant tuberculosis.

**Fig. 8:**
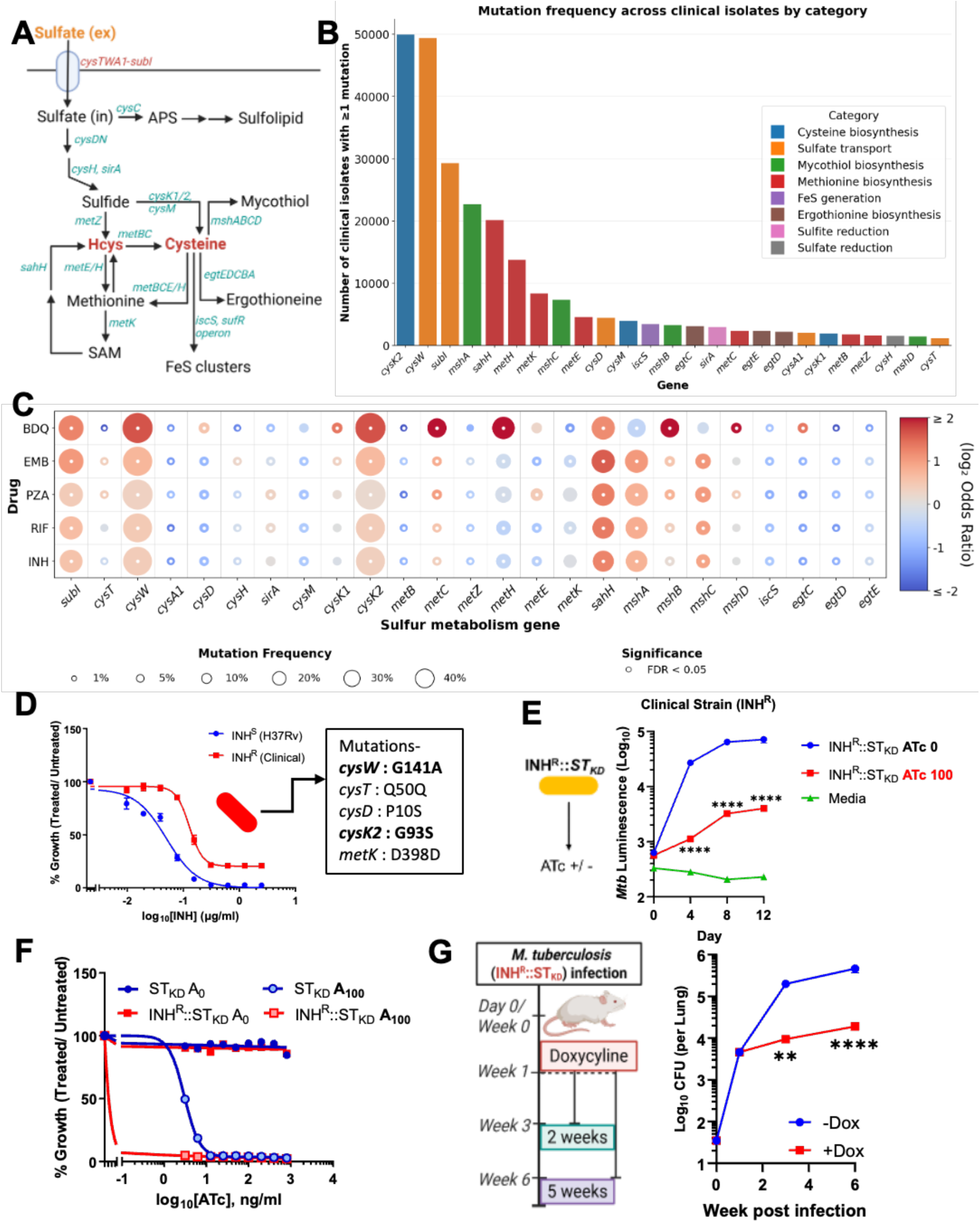
Sulfate uptake inhibition exacerbates hypersusceptibility in clinical drug-resistant *M. tuberculosis*. **(A)** Schematic representation of the key sulfur metabolism pathways and genes involved in the metabolic reactions. (**B**) Mutation frequency in sulfur metabolism genes across clinical isolates in the CRyPTIC database. **(C)** Feature expression heatmap showing the association of sulfur metabolism-related mutations with drug resistance. **(D)** Dose-response curve of H37Rv (INH^S^) and XTB13-198 (BEI Resources, INH^R^) *M. tuberculosis* strains to determine sensitivity against Isoniazid (INH). **(E)** Growth kinetics of a clinically resistant INH^R^ strain with sulfate transporter knocked down using CRISPRi. **(F)** ATc titration of a drug-resistant clinical strain and H37Rv exhibiting sensitivity upon knocking down sulfate transporter(s). **(G)** *In vivo* infection study to determine the fitness of INH^R^ *M. tuberculosis* upon inhibiting sulfate uptake. The data are averages of the three data points indicated by dots and are representative of at least two independent experiments. Error bars correspond to the standard error of the mean (SEM). Statistical significance was assessed by two-way ANOVA (Sidak’s multiple comparisons test) for (E) and (G), where **P = 0.00021 and ****P<0.0001.

## Discussion

The rise of drug resistance necessitates the development of new therapeutic options in the form of novel drugs or treatment strategies. Given the crucial role of sulfur in sustaining life processes, it is an attractive target for combination therapy to enhance the efficacy of existing therapeutic regimens and improve clinical outcomes. In line with this, we elucidated the role of sulfur metabolism in mycobacterial physiology. We primarily used chemically S-depleted (S-free) medium to generate an S-limited transcriptional signature in *M. tuberculosis* and further genetically abrogated the bacteria’s ability to acquire sulfate by knocking down sulfate transporters, thereby expanding our observations in macrophage and murine models of pathogenesis and chemical genetics to elucidate impact of sulfur limitation on bacterial drug susceptibility.

Orthogonally sulfur-starved *M. tuberculosis* (either cultured in S-free medium or genetically knocked down in the sulfate transporter) has reduced intrabacterial sulfur, total thiol content, total protein, and ATP levels; reduced transcriptional-translational activity; and a high NADH/NAD+ ratio, resulting in non-replicating bacilli. This condition is bacteriostatic, as restoration of sulfur supply resumes all stalled physiological processes and bacterial replication. While *M. tuberculosis* can utilise amino acids (cysteine and methionine) and H_2_S as S-sources, it prefers inorganic sulfate as the chief S-source, which is consistent with previous reports (*43*). This suggests that sulfur limitation induces a unique metabolically inactive, reversible state.

Further, S-starved *M. tuberculosis* exhibits major rewiring of core physiological processes, including transcription, translation, proteostasis, oxidative phosphorylation, glycolysis, the TCA cycle, and cell division. Quite similarly, GSMM further highlighted perturbations in metabolic reactions through in silico flux analysis, revealing a signature concordant with the transcriptomic (of S-starved) and proteomic (of ST_KD_) signatures, showing downregulation of several key metabolic reactions involved in the biosynthesis of PDIM, cofactors/ prosthetic groups, pantothenate, CoA, cell-wall biosynthesis, bioenergetics, TCA cycle, and purine and amino acid metabolism.

Sulfur is a constituent of several key cofactors (Thiamine diphosphate, Lipoic acid, Biotin, etc.) and coenzymes (CoSH-A and CoSH-Q) that are required in the enzymatic reactions involved in the catabolism of carbohydrates, sterols, fatty acids, etc., for energy production. Further, reducing equivalents generated as NADH and FADH_2_ transfer electrons to the respiratory chain for ATP biosynthesis via oxidative phosphorylation. Therefore, with the shortage of sulfur, the biosynthesis of these essential components will halt, resulting in nonfunctional enzymes and stalled downstream metabolic pathways. Under such conditions, carbon sources, despite their abundance in the medium, remain unmetabolised, leading to a nutrient-starvation-like situation within the bacterial cell. Downregulation of NDH-1 results in a high NADH/NAD+ ratio and reduced ribosomal assembly and metabolic activity, which are indicative of nutrient starvation (*30*, *44*, *45*). In the event of abundant NADH, cellular priority may shift to rapidly recycling NAD+ for supporting essential survival-related processes rather than maximising proton-pump efficiency. This helps the bacteria maintain a basal physiological state with low energy (ATP) levels through the activity of still functional NDH-2, which is upregulated (*46*). Therefore, one might expect a bactericidal effect upon abrogation of NDH-2 in S-starved bacteria by synthetic lethality (*47*, *48*) – this remains an untested prediction for future work. This is also explained by similar levels of TCA cycle intermediates and amino acids in the *ΔsubI* mutant (which is defective in sulfate uptake) compared to WT *M. tuberculosis* (*43*). This study highlights that sulfur availability influences the carbon and energy metabolism of *M. tuberculosis*.

Sulfur deficiency also increases intrabacterial ROS levels due to reduced thiol content, which can be life-threatening for the pathogen (*43*, *49*). *M. tuberculosis* counters the situation by upregulating redox homeostasis machinery and attempts to recharge the intracellular thiol pool by sulfate acquisition by CysTWA1-SubI and salvaging cysteine and methionine. These observations are somewhat similar to those in S-starved *Pseudomonas aeruginosa* (*50*) and cysteine-deficient *Acinetobacter baumanii* (*49*), where sulfur metabolism-related genes were up-regulated to compensate for the paucity of sulfur in the medium. *M. tuberculosis* attempts to acquire sulfur from the medium via the ABC transporter by upregulating its expression; however, surprisingly, *rv1739c* expression was significantly reduced, indicating that *rv1739c* might not contribute to sulfate uptake, as reported recently (*43*). Interestingly, *rv1066* and *rv1065* were found to be overexpressed and annotated as putative cysteine dioxygenases (CDO) – known for the decomposition of cysteine for sulfur homeostasis in eukaryotes and other bacterial species such as *Bacillus subtilis*, *Bacillus cereus* and *Streptomyces coelicolor* (*51*) - yet uncharacterized in *M. tuberculosis*. These enzymatic reactions can compensate for reduced sulfur content by decomposing cysteine into inorganic sulfate, thereby replenishing the cytosolic thiol pool of the bacterial pathogens – a strategy used by *Pseudomonas aeruginosa* to salvage sulfur from the host during infection (*52*). The existence and operation of such biochemical reactions become increasingly important in the context of host-pathogen interactions.

Furthermore, sulfur limitation reduces bacterial colonisation of macrophages, highlighting the importance of sulfur availability in host-pathogen interactions, as previously reported for *M. tuberculosis* (*43*) and *Salmonella* spp. (*53*). It is also crucial for disease establishment and progression, as evidenced by reduced bacterial burden in infected lungs and improved pathological outcomes in the murine infection model. Despite attenuation, bacterial burden in the knock-down strain continued to increase over time, indicating that *M. tuberculosis* can regenerate its thiol pool by salvaging S-containing metabolites, such as cysteine and methionine, from surrounding host tissues. In addition, host-generated H_2_S supports bacterial growth and survival, as previously reported and extensively reviewed (*16*, *17*, *37*, *53*). Indeed, co-inhibition of host H_2_S biosynthesis and bacterial sulfate acquisition exhibited severe attenuation in macrophages, supporting host-directed inhibition of H_2_S biosynthesis as a complementary strategy to a sulfur transporter-targeted therapeutic.

Chemical genetics under in vitro as well as in vivo settings revealed that restricting sulfur uptake significantly predisposes *M. tuberculosis* to specific anti-TB drugs. This vulnerability manifests as pronounced shifts in MICs against respiration- and growth-related antibiotics, specifically RIF, BDQ, EMB, and CFZ. The enhanced efficacy of respiration-related antibiotics (BDQ and CFZ) may relate to disruption of the respiratory chain containing FeS cluster-containing complexes; this mechanism was not directly tested in this study. In contrast, sulfur limitation does not impact the effectiveness of INH. This may be attributed to its mechanism of action, which selectively targets replicating rather than non-replicating bacteria. Furthermore, an elevated NADH/NAD+ ratio in S-starved *M. tuberculosis* counteracts INH activity (*54*, *55*). Drug tolerance is a concern in the context of antibiotic therapy, as bacteria gain phenotypic ability to withstand drug treatment for longer periods, facilitating the development of resistance (*56*). Treating sulfur-deprived bacteria with higher antibiotic concentrations reduced the number of tolerant bacteria. Comparatively, in contrast to INH, drugs targeting transcription (RIF) and respiration (BDQ) effectively reduced tolerance in S-deprived bacteria. We note that sulfate transporter knockdown alone is bacteriostatic rather than bactericidal and that the synthetic lethal phenotype we describe emerges specifically upon combination with antibiotic treatment. This is in concert with classical synthetic lethality applied here to chemical-genetic rather than purely genetic-genetic interaction. These findings emphasise that in addition to potentiating the cidal activity of anti-TB drugs, the potential of sulfur limitation can be harnessed to address drug tolerance in *M. tuberculosis*.

Interestingly, animal model studies revealed distinct drug-sensitivity dynamics. INH treatment of *M. tuberculosis* defective in sulfur acquisition sterilised the infected mouse lungs within 4 weeks of treatment. These observations highlight the complex dynamics among growth, metabolism, and drug efficacy. During the early phase of infection, the mutant was unable to take up inorganic sulfur, thereby affecting bacterial growth within the host. Over time, as part of an innate immune response, the host generates H_2_S, which the pathogen actively scavenges along with other thiol-containing metabolites, replenishing its thiol pool and thereby restoring growth, supporting a model in which sulfate transporter-lacking *M. tuberculosis* gains a steady rise in CFU despite marked attenuation. Interestingly, this transition back to active growth increased the pathogen’s vulnerability to INH, leading to sterilisation of the infected lung tissues in mice. Consistent with H_37_Rv, abrogation of sulfur uptake in clinical drug-resistant (INH^R^) *M. tuberculosis* slowed its growth and rendered it hypersensitive to even low-level sulfur depletion *in vitro*. Similarly, its replication was curtailed, effectively controlling the bacterial burden in the lung tissues of infected mice. Mutations in the sulfur metabolism-related genes (amongst other mutations) may have led to a constrained thiol pool in the clinical strains, resulting in slow growth. Further restricting access to sulfur in such strains using CRISPRi led to pronounced fitness cost. Likewise, rifampicin-resistant clinical isolates were shown to harbour mutations in sulfur acquisition and assimilation genes, rendering them slow-growing (*57*). In sum, these findings demonstrate that sulfur metabolism remains essential for the survival of *M. tuberculosis*, regardless of its genetic background or drug susceptibility profile.

Despite several promising findings, there are a few caveats to the present study. Indeed, non-synonymous genetic mutations (SNPs) are associated with *loss-of-function* phenotypes; however, most of the time, such mutations can be well tolerated in clinical isolates and must therefore be validated experimentally. Further, genetic association analysis was not designed to control for phylogenetic lineage or clonal background; mutations in sulfur metabolism genes were observed across both drug-resistant and drug-sensitive isolates and across the majority of represented lineages, and the associations reported here should be interpreted as descriptive rather than causal.

Collectively, this study demonstrates that sulfur acquisition is important for bacterial growth and metabolism and that it also contributes to pathogenesis and the ability to establish infection. A reduction in intracellular sulfur levels decreases the thiol pool, making *M. tuberculosis* more sensitive to host-generated assaults, such as ROS and RNS, that restrict bacterial replication. The absence of sulfur rewires bacterial metabolism, thereby sensitising them to anti-tubercular drugs. This approach, involving pharmacological inhibitors of sulfate transporters, can be effectively harnessed to control mycobacterial infection, including drug-resistant cases, either alone or in combination with other antibiotics **(Fig. 9)**. Furthermore, the fact that host cells lack an ABC-sulfate importer presents an opportunity to target the bacterial sulfate uptake system as a potential candidate for drug discovery and development.

**Fig. 9:**
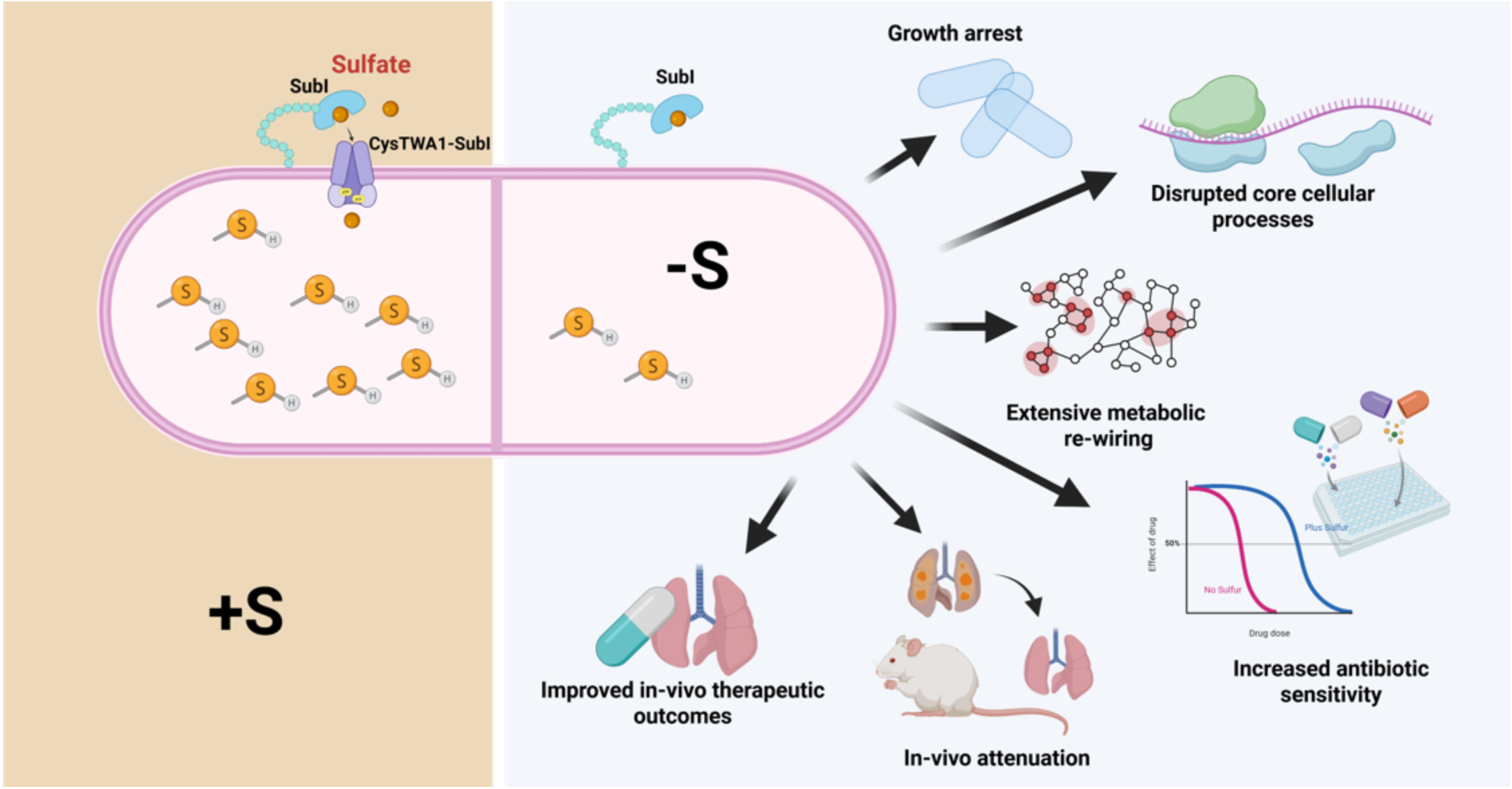
**Graphical abstract depicting consequences of inhibiting the acquisition of sulfur uptake in *M. tuberculosis***. *M. tuberculosis* acquires inorganic sulfur in the form of sulfate/ sulfites, etc., which is crucial for maintaining the thiol pool of the cells that protects bacteria from ROS/ RNS assaults imposed by the host. Upon inhibition of sulfur acquisition, the thiol pool is drastically reduced, triggering extensive metabolic rewiring that slows metabolism, alters respiration, halts transcription and translation, resulting in growth arrest and reduced biomass. Sulfur limitation renders *M. tuberculosis* sensitive to existing anti-TB drugs, thereby improving their efficacy. Sulfur limitation also affects the pathogenicity of *M. tuberculosis* in vivo and results in sterility upon treatment in combination with INH.

## Materials and Methods

### Bacterial strains and culture

All bacterial strains were cultured in 7H9 medium (Difco, BBL) supplemented with OADS (Oleic acid, Dextrose, and NaCl, i.e., Salt) and 0.01% Tween 80. Bacterial enumeration by CFU was performed on 7H11 agar plates supplemented with OADS, as mentioned, and colonies were counted after 21 days of incubation. Kanamycin (Sigma), 35 μg/ml; Hygromycin (Sigma), 50 μg/ml; and Zeocin (Invitrogen), 25 μg/ml were used as required. Phosphate Buffered Saline (PBS) was used at a final concentration of 1X wherever mentioned, with or without 0.01% Tween (PBST/ PBS, respectively). Salts used for the preparation of minimal medium were purchased from Sigma unless mentioned otherwise. Anhydrotetracycline chloride (ATc, Cayman) was used at final concentration 100ng/ml unless specified (*58*).

### Preparation of Sulfur-free (S-free) medium

The sulfur-free medium was prepared by replacing the sulfate-containing salts (MgSO_4_ and ZnSO_4_) with chloride salts (MgCl_2_ and ZnCl_2_) in the minimal medium as described elsewhere (*59*). The rest of the composition was the same with different carbon sources when desired – glycerol (0.2% w/v), cholesterol (200µM) or sodium propionate (10mM).

### CRISPRi-mediated gene knockdown

Plasmid constructs harbouring target gene knockdown cassettes for individual genes – *cysT* (*Rv2399c*), *cysW* (*Rv2398c*) and *sulP* (*Rv1739c*) were generated by cloning respective guide oligo pairs in the BsmBI-digested vector backbone plJR965 (*33*). Next, the *cysT*-targeting construct was digested with SapI, and the genetic cassettes targeting *cysW* and *sulP* were PCR-amplified using primers with compatible overhangs via Golden Gate cloning (*34*). The resulting plasmid construct targeting all three genes, i.e., *cysT*, *cysW*, and *sulP*, in tandem was confirmed by sequencing. The triple-knockdown construct was transformed into *M. tuberculosis:lux* (bioluminescent reporter strain, luciferase-expressing plasmid (*60*)) by electroporation and recovered on kanamycin/zeocin-supplemented 7H11 agar plates. A few colonies were screened for a functional knockdown system by ATc titration.

### ATc titration assay for defining vulnerability

The ATc titration assay was performed in 7H9-enriched medium as described previously (*61*). Triple-knockdown and non-targeting (NT) control strains were inoculated at OD 0.005 and exposed to 10-point, 2-fold dilutions of ATc (Cayman), starting from 1600 ng/ml. The plates were incubated for 72 hours at 37°C without shaking. Auto-bioluminescence was measured using a plate reader (SpectraMax ID3, Molecular Devices), and per cent growth was calculated relative to ATc-untreated cultures.

### RT-qPCR for knockdown confirmation

Total RNA was extracted from log-phase cultures of *M. tuberculosis*, either treated or untreated with ATc (100 ng/ml) to knockdown the target genes. Cells equivalent to OD 2 (∼6x10^8^ bacteria) were pelleted down, washed with PBST and resuspended in 1ml RNAIso plus (TaKaRa) followed by cell lysis by bead beating. 0.2 ml of chloroform was added, and the samples were mixed by agitation and centrifuged (12000 rpm, 10 min at 4°C) to facilitate phase separation. The aqueous phase was collected, and RNA was precipitated with isopropanol, followed by two washes with chilled 70% ethanol. RNA was air-dried and resuspended in nuclease-free water (Invitrogen, Ambion). Genomic DNA was removed by TURBO DNase treatment (Invitrogen, Ambion). cDNA synthesis was performed using 5 µg DNase-treated RNA and random hexamers via reverse transcription. Next, target genes’ expression was determined by qPCR on a QuantStudio6 system (Thermo Scientific) using gene-specific qPCR primers. Knockdown was quantified by the ΔΔCt method (*62*), with ATc 0 as the normalisation counterpart, and *sigA* as the housekeeping control.

### Growth kinetics

Growth kinetics for the bacterial strains were performed as detailed elsewhere (*58*). Bacterial viability was determined by measuring optical density (OD), luminescence, and CFU at the indicated time points, as required. For sulfur-related growth kinetics of *M. tuberculosis* H37Rv, the bioluminescence reporter strain was grown to mid-log phase (OD 0.4-0.5), washed with PBST and inoculated at an OD of 0.05 in sulfur-rich and sulfur-free media. 100 µl aliquots were drawn from the culture into a 96-well plate (white, flat-bottom, SPL), and luminescence was recorded at the indicated time points using a Multimode Microplate reader (SpectraMax Pro ID3, Molecular Devices). Additionally, OD of the cultures was measured, and appropriately diluted samples were plated over 7H11 agar plates, followed by incubation for 20-21 days and enumeration of colonies.

For sulfur re-supplementation growth kinetics, different S-containing sources - MgSO_4_, (NH_4_)_2_SO_4_, cysteine (100µM) and methionine (50µg/ml) were added to individual bacterial cultures growing in Sulfur-free medium.

For growth kinetics of CRISPRi strains, cultures were back-diluted to OD 0.01 in fresh medium and supplemented with ATc to initiate CRISPRi knockdown and sulfur sources – cysteine (100 μM), methionine (50 μg/ml) and NaHS (200 μM) whenever desirable.

### Sulfur estimation by Inductively Coupled Plasma-Mass Spectrometry (ICP-MS)

Bacterial cultures were grown to an OD of 0.4-0.5, washed with PBS, and diluted back to an OD of 0.05 in both sulfur-rich and sulfur-free media, then allowed to grow for 5 days. After 5 days, samples were processed as described elsewhere (*58*, *63*). Briefly, after the incubation period, the cells were washed twice with sterile deionised water (ICP-MS grade) to remove residual medium and salts. Aliquots of cells (OD 1.0) were treated with 70% conc. HNO_3_ (Trace metal analysis-LCMS grade) to digest the cellular biomass. 100 µl of 30% H_2_O_2_ was added to the acid-treated samples for microwave digestion with a ramp of 250W for 10 minutes and a power hold of 250W for 10 minutes. Digested samples were then diluted (1:100) using deionised water. Next, these diluted samples were analysed using ICP-MS (iCAP TQ, Thermo Fisher Scientific). Mg standard was run at concentrations (1 ppb to 1000 ppb), while a ^34^S standard was run at concentrations (0.1 ppb to 1000 ppb) to prepare the calibration curve. ICP-grade water was used for standard preparation, sample dilution and as a blank. Samples were run with a dwell time of 0.1 sec, and 1% HNO_3_ was run for 30 sec between samples to remove carryover from the previous sample. The average value of three runs was used to calculate the magnesium and sulfur concentrations in each sample. All the samples were run in KED mode (Kinetic energy dissociation) to remove polyatomic interference. Qtegra software (Thermo Scientific) was used to operate the instrument. Blank-corrected values were plotted against the respective samples.

### Gene Expression Analysis of S-starved *M. tuberculosis*

For sample preparation, *M. tuberculosis* H_37_Rv (WT) culture was grown to the early log phase, and cells were harvested when the OD reached 0.5. The cells were washed with PBST, transferred to sulfur-rich and sulfur-free media at an OD of 0.5, and cultured in triplicate per group. Post 5 days of incubation, total RNA was isolated using the Trizol method as described. Cultures were pelleted, washed twice with PBS and cell pellets were processed for total RNA isolation by Trizol method. Extracted RNA was subjected to DNA removal using TURBO DNaseI. NEB Ultra II directional RNA-sequencing Library Prep kit protocol was used to prepare libraries for total RNA-Seq (NEB). An initial total RNA concentration of 500ng was used for cDNA synthesis and library preparation. Initially, the raw reads were processed using *fastp* and mapped to the reference genome using *bowtie2*, and genes with low expression values (mean FPKM value <0.5) were treated as background (i.e. ‘noise’) and hence excluded from further analysis. After noise removal, the gene expression values for each experimental condition were merged, and the final data were analysed. To assess the variation in the gene expression data, we performed principal component analysis (PCA) using the *prcomp* package in “R”, which applies the singular value decomposition to the data matrix. For the comparison between gene expression profiles from S-free and S-rich bacterial cultures of *M. tuberculosis*, genes with a fold change of 2 (log_2_FC > ± 1) and false discovery rate (FDR)-adjusted p-value ≤ 0.05 were considered differentially expressed. The Benjamini-Hochberg multiple hypothesis testing was applied to obtain the FDR-adjusted *p*-value using the *DESeq2* package in “R” (*64*). Subsequently, volcano plot and feature expression heatmaps were generated in “R” using *ggplot2*, *enrichplot* and *gplots* packages.

### Genome-Scale Metabolic Modelling

#### Data pre-processing

The high-throughput RNA-seq dataset underwent initial preprocessing, which involved removing genes expressed in fewer than 50% of replicates in each condition. Outliers were identified using the interquartile range (IQR), defined as IQR = Q3-Q1, where Q1 and Q3 represent first and third quartiles, respectively. For each gene in each condition, replicates with expression values below the lower bound (Q1-1.5*IQR) or above the upper bound (Q3+1.5*IQR) were considered outliers. To correct these, the counts were replaced using the minimum (m) and maximum (M) of the non-outlier replicate expression values for that gene in the corresponding condition. Values below the lower bound were replaced with ‘m’, whereas values above the upper bound were replaced with ‘M’. Finally, the expression values for each gene were normalised by dividing by the maximum expression count of that gene across all replicates within the same condition. Replicate expression counts within each condition were then aggregated by computing their mean.

#### Mapping gene expression to reaction activity scores using GPR rules

The gene-protein-reaction (GPR) rule assigned to each reaction in the model provides information about the genes required to catalyze that reaction. These rules use logical statements to describe how genes combine to form functional enzymes. AND indicates genes that function together as part of a protein complex, while OR represents alternative genes that can catalyze the same reaction. For each reaction, the corresponding GPR rule was first split into its OR components, each representing an alternative enzyme capable of catalyzing the reaction. Within each OR block, the presence of one or more AND operators were evaluated. When no AND operator was present, the reaction was associated with a single gene, and the gene’s expression value was used directly. When an AND relationship was encountered, indicating that multiple genes form a single enzyme complex, the expression levels of all participating genes were collected, and the complex activity was defined as the minimum of these values. After processing each of the OR groups for a reaction, the resulting contributions were combined to determine the overall reaction-level expression. Following established E-Flux methodology, OR relationships were treated as additive; therefore, the activity scores from all OR blocks were summed to obtain the final reaction-level expression value. This procedure was performed for every reaction in the model and applied independently to each experimental condition.

#### Building context-specific metabolic models

Context-specific metabolic models were constructed using the E-Flux method (*26*). This method uses gene expression-derived reaction activity scores to constrain reaction flux bounds. Reactions catalyzed by highly expressed genes were allowed to carry higher fluxes, whereas reactions associated with low gene expression were more tightly constrained. Thus, the steady-state solution space is described as:

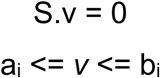

where *v* is a flux vector, S is the stoichiometric matrix, and *a_i_* and *b_i_* represent minimum and maximum fluxes through the reaction *r_i_*.

For reversible reaction: *a_i_* = *-b_i_*, where *b_i_* = f (reaction activity score of *r_i_*)

For irreversible reaction: *a_i_* = 0

#### Mean match flux evaluation

Flux sampling was performed using the GP sampler, a uniform random sampling algorithm implemented in the Constraint-Based Reconstruction and Analysis (COBRA) Toolbox (*65*). The GP sampler is derived from and improves upon the Artificial Centring Hit-and-Run (ACHR) algorithm and operates by iteratively selecting a random direction and step size, ensuring each new point remains within the feasible, linearly constrained solution space (*27*, *28*, *66*). The algorithm samples the flux space with a fixed number of points, with a default sample size of approximately twice the number of reactions in the model. In our models, this default setting produced 2,476 sampled feasible flux vectors. To obtain a representative steady-state flux distribution for this sampling size, we defined the solution as the mean flux vector of all sampled points. To assess the robustness of the mean flux vector, we increased the number of sampled flux vectors to 10-fold and 25-fold of the default sample size for the control (S-enriched) and test (S-deprived) conditions, respectively. For each sample size, we computed the mean flux vector and assessed convergence by computing the Euclidean distance between mean vectors from successive simulations. The sampling procedure was continued until the distance between the mean flux vectors stabilised at a minimum, indicating convergence (*67*).

### Total Protein Estimation

The bacterial culture was grown to an OD of 0.4-0.5 (early log phase) and harvested by centrifugation at 4000 rpm for 10 minutes at RT. The cells were washed with PBST and transferred to minimal salts medium, either or not supplemented with sulfur-containing salts, at an OD of 0.05. Post 5 days of incubation, OD 1.0 equivalent cells were washed with PBS and resuspended in 1000µl PBS (with PIC) followed by lysis using bead beating method with 0.2µ diameter zirconia beads for 5 cycles of 45 seconds with 1 minute of intermittent cooling on ice. The lysates were clarified by centrifugation at 13000 rpm for 10 minutes at 4°C, and the supernatants were collected in fresh MCTs. The clarified lysates were filtered through 0.22 µm filters to remove any remaining bacterial cell debris. The protein content of the lysates was measured using the BCA protein estimation kit according to the manufacturer’s instructions. Total protein content (mg/ml) was normalised to the OD of the cells used for lysis.

### ATP Estimation

Bacterial cultures were grown to an OD of 0.4-0.5 (early log phase), and cells were harvested by centrifugation at 4000 rpm for 10 minutes at RT, followed by washing with PBST. The cells were back-diluted to an OD of 0.01 in S-rich or S-free (or ATc 100ng/ml for CRISPRi strains) media. After 5 days of incubation, the cells were washed with PBS, and OD 1.0 equivalent cells were resuspended in 1ml PBS (with PIC), followed by lysis by bead beating with 0.2 µ-diameter zirconia beads for 5 cycles of 45 seconds with 1 minute of intermittent cooling on ice. The lysates were clarified by centrifugation at 13000 rpm for 10minutes at 4°C, and the supernatants were collected in fresh MCTs. The clarified lysates were filtered through 0.22 µ filters to remove any remaining bacterial cell debris. 50μL of clarified lysates was transferred to a 96-well plate (white, flat-bottom, SPL) and mixed (1:1) with BacTiter-Glo (Promega) reaction buffer, followed by incubation at 37°C for 30 minutes. PBS was used as a blank to cancel out the background signals from the samples. Luminescence was recorded using a Multimode Microplate reader (SpectraMax Pro ID3, Molecular Devices). Background-corrected ATP levels were expressed as relative units relative to the S-rich cells.

### Thiol Estimation

Bacterial cultures were grown to log phase in the respective media (with or without sulfate or ATc 100ng/ml for CRISPRi strains) and washed with PBST. Cells equivalent to OD 1 were lysed by bead-beating (5 cycles, 45 seconds with 1-minute intermittent cooling) and filtered through 0.2μm pore-size filters. 100 μL of freshly prepared 2X Thiol assay buffer (1 M Tris, pH 7.0, 200 μM DTNB) was added to wells in a 96-well plate containing 100 μL of cell lysates. The plate was incubated for 20 mins at room temperature, and a colorimetric readout at 410nm was performed using a microplate reader (SpectraMax ID3, Molecular Devices). Thiol content was normalised by total protein across all the samples.

### Bacterial replication rate estimation using the Replication Clock plasmid

*M. tuberculosis* H37Rv (WT) bioluminescence reporter strain carrying mycobacterial replication clock plasmid-*pBP10*, Kan^r^ (*21*, *68*) was grown to an OD of 0.4 - 0.5. The cells were harvested and washed with PBS to remove residual sulfur from the previous medium. The cells were then inoculated into sulfur-rich and Sulfur-free media at an OD of 0.05. 100 µl aliquots from the cultures were serially diluted and plated on 7H11 plates supplemented with or without Kanamycin. Plates were incubated for 20-21 days until the appearance of colonies which were scored and used to calculate %pBP10+ cells (cells retaining replication clock plasmid) as per the expression below:

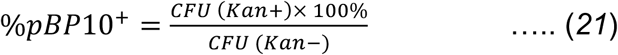

Further, the other factors – replication rate, net growth rate, plasmid loss rate, death rate, etc. were determined as described by Gill *et. al.* (*21*) using a custom-written script with the scripting language “R”.

### Intrabacterial NADH/NAD+ assay using ratiometric biosensor Peredox

Mycobacterial strains harbouring plasmid pIMT100 (pMV762:Peredox-mCherry) were grown in respective media (with or without sulfur or ATc 100 and 500ng/ml for CRISPRi) until log phase, and cells were harvested by pelleting down (4000rpm, 10mins, RT). Pellets were washed and resuspended in PBST at an OD of 0.2. 100 μL of cell suspensions were aliquoted into a 96-well plate (black, flat-bottom), and fluorescence was recorded at wavelengths 510 and 615 nm by exciting the samples at 400 nm and 587 nm for cpmTS and mCherry, respectively, using a microplate reader (SpectraMax ID3, Molecular Devices). The fluorescence readouts were background corrected, and the ratios of fluorescence emissions at 510 nm to 615 nm (green/red) were plotted for all samples to estimate the intrabacterial NADH/NAD+ ratio in bacterial cells exposed to different conditions at the indicated time points.

### Intrabacterial Sulfate assay using ratiometric biosensor mThyone

Mycobacterial cultures expressing the mThyone biosensor (from episomal plasmid pST-H:*mThyone*) were cultured under respective culture conditions (with or without sulfur or ATc 100 and 500ng/ml for CRISPRi), harvested, and washed with PBST. Pellets were resuspended in PBST at a density of OD 0.2, and 100 μL of the respective samples were aliquoted into a 96-well plate (black, flat-bottom). Fluorescence was determined by exciting the samples at 485 nm and 555 nm while recording the fluorescence emission at 530nm and 595 nm (for GFP and RFP, respectively) using a microplate reader (SpectraMax ID3, Molecular Devices). The fluorescence signals were background corrected, and the ratio (Green/Red) was determined for all samples, indicating sulfate levels in the bacterial cells.

### Single Cell Fluorescence Dilution (SCFD) assay

*M. tuberculosis* dual reporter strain carrying an episomally replicating plasmid with mEmerald (GFP) under constitutive promoter (*P_hsp60_*) and Tet-inducible tagRFP under *P_uv15tetO_* promoter (*69*) was “pre-loaded” with RFP by ATc (500ng/ml) induction (*70*) until the culture attained an OD of 0.5-0.6. Next, bacterial cells at an OD of 0.2 were inoculated into 2ml S-free medium and cultured for 4 days. After this, ammonium sulfate or GYY4137 (H_2_S donor) was added to the designated groups, and fluorescence dilution was allowed to occur. Individual cultures per group at each time point were pelleted down, washed once with PBST and fixed with 4% paraformaldehyde (PFA), followed by flow cytometric analysis as described previously (*70*, *71*). For analysis, the GFP+ singlets were distinguished from debris and cells with RFP signals (low and high) were analysed for dilution over time. For determining the dilution of RFP over time, the following relation was used:

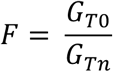

where *F* is the fluorescence dilution factor, G_T0_ is the RFP fluorescence geometric mean of the population at T0, and *G_Tn_* is the fluorescence geometric mean of the growing population at *T_n_* (*71*).

Further, the RFP gets diluted (halved) per cell division; therefore, the number of generations, *N,* at a specific time, was calculated as:

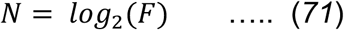

### Transcription-Translational Activity (TTA) Assay

TTA was performed using a dual-reporter *M. tuberculosis* strain as described elsewhere (*72*) with some modifications. Briefly, the dual-reporter strain was grown to log phase, and harvested cells were washed and diluted back to an OD of 0.05 in 10mL of S-rich and S-free media. ATc (500 ng/ml) was added to the cultures at the start of the experiment to initiate transcriptional induction of Tet-controlled RFP in all groups. On the 8th day, ammonium sulfate was added to a separate group cultured in S-free medium to determine time-period-specific transcriptional activity rescue in the supplemented cultures. 100 μL of the bacterial cultures were aliquoted into a 96-well plate (black, flat-bottom), and fluorescence intensities with excitation at 485nm and 585nm, and emission at 525nm and 625nm for mEmerald and tagRFP, respectively, were recorded using a microplate reader (SpectraMax ID3, Molecular Devices). The fluorescence readouts were background corrected and analysed. While GFP signals monitored bacterial growth across groups, RFP signals revealed the ability of cells in each group to transcribe and translate RFP in response to ATc induction.

### Proteomics by Label-free mass spectrometry

#### Sample preparation

*M. tuberculosis* strains - Non-targeting (NT) and sulfate transporter(s) (ST_KD_) were grown in the presence of ATc (100ng/ml), and cells equivalent to an OD of 1 were pelleted. Pellets were washed with PBST followed by lysis in PBST + PIC by bead beating. The lysates were clarified by centrifugation at 12000rpm for 10 min at 4°C and filtered through 0.2 μm pore-sized filters. The filtered lysates were then dried by SpeedVac and processed for mass spectrometry.

#### Sample processing

In-sol trypsin digestion was carried out as described earlier (*73*) with some modifications. Briefly, 50µg protein was denatured by 8M urea at 1mg/ml concentration. 10mM DTT was used for disulfide bond reduction for 1 h at 37°C, followed by alkylation with 20mM IAA in the dark. 50 mM ammonium bicarbonate was added to the sample mixture to dilute the urea concentration to 2 M. Proteins were digested with trypsin at 1:20 for overnight at 37°C. Following digestion, the resulting peptide mixture was acidified with formic acid, desalted, and vacuum-concentrated.

#### Mass Spectrometry conditions

LC-MS/MS was performed using a Vanquish™ Neo UHPLC System, connected on-line with an Orbitrap Exploris™ 480 Mass Spectrometer (Thermo Scientific). The trypsin-digested samples were desalted offline by a C18 column prior to loading them on a Trap (PepMap™ Neo Trap Cartridge 100 A, 5 μm C18 0.3 mm x 5 mm) and an analytical nano column (Easy-Spray™ PepMap™ Neo 2 μm C18 75 μm x 150 mm). The mobile phase for HPLC was as follows: water/formic acid (100/0.1%) and acetonitrile/formic acid (80/0.1%). A sample (500 ng) was injected on the column with a flow rate of 300 nl/min. Analytical separation was established using the following gradient conditions: initial 6% B for 2min, then a linear gradient to 31% B over 58 min, followed by another linear gradient to 44% B over 30 min. Following the peptide elution window, the gradient was increased to 100% B in 1 min and held for 9 min.

#### Data Acquisition

A data-dependent acquisition experiment was performed on an Orbitrap Exploris™ 480 system having a FAIMS Pro™ Interface. FlexMix calibration solution (Pierce) was used to tune the instrument before data acquisition. The sample data were acquired using a nano-spray ionisation ion source with a voltage of 2000 V, ion transfer tube temperature of 280°C, FAIMS voltage of -50 V, RF lens of 40%, automatic gain control of 300%, maximum injection time in auto mode, and 1 microscan. The MS was operated at a resolution of 60,000 FWHM and a mass range of m/z 350-1200. A total of 30 survey scans were acquired at 15,000 resolution and 28% HCD collision energy if exceeding an intensity threshold of 5.0e3 with a 2+ to 6+ charge states. The dynamic exclusion duration was set to 45s.

#### Database search for protein identification

Identification of proteins in the samples was performed via database searching against the *M. tuberculosis* H37Rv, obtained from the UniProt database with Proteome Discoverer software having *Chimerys* and *INFERSYS* rescoring algorithms, using the following search parameters: enzyme trypsin; maximum missed cleavage 2; mass tolerance 10 ppm; static modification Cys carbamidomethyl; dynamic modification Met oxidation; target/decoy strategy using Percolator; target FDR 1%.

#### Analysis

A custom script was written using the language “R” to analyse the acquired proteomics data. To begin with, proteins having protein abundance values in at least two of the three replicates in both the control (NT) and test (ST_KD_) groups were selected for analysis. Protein abundance values were log-transformed, normalised, and then imputed. Imputed data were used to calculate fold changes, and Benjamini-Hochberg multiple hypothesis testing was applied to obtain FDR-adjusted *p*-values. Differentially expressed proteins (DEPs) were further used for pathway analysis, and representative proteins were depicted in heatmaps prepared using *TBtoolsII* (*74*).

### Mutation Analysis in Clinical Isolates

To investigate if the clinical isolates have compromised sulfur metabolism, we used the gene variation datasets from the CRyPTIC database (*40*, *41*). The CRyPTIC database (version 3.4.0) consists of whole-genome sequences (WGS) and phenotypic Drug Susceptibility (pDST) data for 53,897 clinical isolates, while only pDST data are available for an additional 11,595 samples (*41*). From the CRyPTIC datasets, mutations (SNPs, insertions, deletions, frameshifts, stop gained, etc.) in the genes involved in sulfur metabolism were quantified across all clinical isolates, including drug-resistant and sensitive strains. Further, the frequency of synonymous and non-synonymous mutations per gene in all isolates was determined, and the association of these mutations with drug resistance was assessed by Fisher’s exact test with no lineage adjustment, since we intended to examine genetic mutations irrespective of the lineages to which these isolates belong. Finally, the frequencies of individual mutations per gene involved in translating to a faulty protein (insertions, deletions, non-synonymous mutations, etc.) were identified against their drug-resistance backgrounds.

For XTB13-198, a clinical isolate obtained from BEI resources, the WGS raw data (SRR1172224 and SRR1168991) (*42*) were downloaded from NCBI using the *prefetch* function of the SRA toolkit and converted to paired-end fastq files with the *fasterq-dump* tool. Sequencing read quality was checked using *fastqc,* followed by adapter removal using *fastp*. Alignment to the reference genome was performed by BWA and SAMtools. BCFtools was used for variant calling with ploidy 1 and optimised filters: QUAL<30, DP<10, MQ<40 and AF<0.75 to eliminate low-quality reads. A custom Python script was used for variant annotation to identify mutations.

### Minimum Inhibitory Concentration (MIC) assay

To determine the MICs of the drugs upon knocking down the sulfate transporter in Bioluminescent *M. tuberculosis*, the cultures (NT and ST_KD_) were grown to log phase, diluted back to 0.01, and exposed to 10-point, 2-fold dilutions of the indicated antibiotics – Rifampicin, Isoniazid, Bedaquiline, Ethambutol and Clofazimine. At day 4, luminescence was recorded, and MIC was calculated by non-linear regression using the GraphPad Prism as described previously (*75*).

### *In-vitro* drug efficacy assay

*M. tuberculosis* strains depleted for genes of interest using CRISPRi were prepared by pre-treating them with 100ng/ml ATc. The ATc pre-treated cultures depleted of the target genes were diluted back into fresh 2ml 7H9 medium to an OD of 0.2 and exposed to different concentrations of ATc (0, 100, 500, 1000 ng/ml) only or higher MIC concentrations of antibiotics Rifampicin (10X), Isoniazid (10X), and Bedaquiline(5X) with ATc (100 ng/ml). At day 7 post-treatment, aliquots of cultures were appropriately diluted and plated on 7H11 agar plates, then incubated for 18-21 days, after which colonies were scored, and survival was determined.

### Macrophage infection

Mycobacterial CRISPRi strains were either or not pretreated with ATc (100 ng/ml) for 4 days prior to infection. THP1-derived macrophages were seeded at a density of 5×10^5^ cells/well in a 24-well plate and differentiated with PMA (Sigma). Cells were maintained in RPMI medium supplemented with 10% fetal bovine serum. Next, PMA was removed 24 hours post-seeding, followed by a 24-hour rest. An MOI of 1 was used to infect macrophages with mycobacterial strains for 4 hours, followed by three washes with pre-warmed PBS. To assess bacterial uptake, ATc (100 ng/ml) was added to the infection medium. The macrophages were lysed with lysis buffer (PBS + 0.01% Triton X-100), and serially diluted samples were plated on 7H11 agar plates. Respective inocula were plated to determine the input bacterial density for each strain. For 96-well plate-based experiments, macrophages were seeded at a density of 5×10^4^ cells/well, and infection was performed with bioluminescent CRISPRi strains (NT::lux and ST_KD_::lux) at an MOI of 2 (*60*); ATc was added to initiate knockdown of target genes. D,L-Propargylglycine (PAG) was added to the respective wells at a concentration of 10 μM. Intracellular growth was monitored either by recording luminescence or CFU plating.

### Animal infection

Sulfate transporter knockdown *M. tuberculosis* strain (ST_KD_) was grown to log phase, de-clumped by passing through an 18-gauge syringe 5-6 times, and a suspension of 3×10^8^ cells/ml was prepared. BALB/c mice (female) were infected by aerosolisation with the single cell suspension, and a few representative mice (n=5) were sacrificed to assess bacterial deposition. At day 1 post-infection, infected mice were randomised into three groups and doxycycline (2mg/ml in 5% sucrose) was administered in drinking water to induce CRISPRi of the transporter genes. This group was marked as “+Dox (W0)”. Similarly, another group received doxycycline treatment at week 1 post-infection and was marked as “+Dox (W1)”. A control group that received only 5% sucrose drinking water was designated “-Dox” and was isogenic to wild-type *M. tuberculosis* with a functional sulfate acquisition system. At indicated time points, animals (n=5 per group) were sacrificed, and the organs (lungs and spleen) were harvested and homogenised. The homogenates were serially diluted and plated over 7H11 agar plates, followed by CFU enumeration. At week 6 post-infection, portions of the tissue were preserved in formalin and processed for H&E staining to assess tissue damage, including granulomatous lesions, in the Dox-treated (W0) and untreated groups. The experiment was conducted in accordance with the guidelines of the CCSEA, Government of India, with approval from the Institutional Animal Ethics Committee (IAEC), BRIC-THSTI, bearing number IAEC/THSTI/210.

Similarly, the Clinical INH^R^ *M. tuberculosis*-derived ST_KD_ strain was used to infect BALB/c mice, and one-week post-infection, doxycycline (2mg/ml in 5% sucrose) was administered via drinking water, with only sucrose in the control group. Animals (n=5) were sacrificed, and lungs were harvested, homogenised, and plated over 7H11 agar plates. Bacterial burden was assessed by CFU enumeration post 20-25 days of incubation. The experiment was conducted with approval from the IAEC, BRIC-THSTI, under approval number IAEC/THSTI/258.

### Drug efficacy in the animal model

*M. tuberculosis-*derived ST_KD_ strain was grown and administered to BALB/c mice through the aerosol route. On day 1 post-infection, mice (n=5) were sacrificed to assess baseline infection. The mice were randomised into two groups: one received doxycycline (2mg/ml in 5% sucrose) to activate CRISPRi, while the other received only a 5% sucrose (w/v) solution as a control. The disease was allowed to progress through week 2 when Isoniazid (10mg/kg body weight) was administered through oral gavage 5 days/week for 4 weeks. A separate Untreated (UT) control was maintained for both -Dox and +Dox groups. Animals (n=5) from each group were sacrificed, and the lungs were harvested, homogenised, and plated on 7H11 plates to determine bacterial burden post-drug treatment. The animal experiment was conducted in accordance with guidelines laid down by the CCSEA, Government of India, and was approved by the IAEC, BRIC-THSTI, with protocol number IAEC/THSTI/365.

## Data Availability

1. Raw data for transcriptomics of S-starved *M. tuberculosis* is available at the Indian Nucleotide Data Archive (INDA), IBDC, India. Accession ID: PRJEB64040 (https://ibdc.dbt.gov.in/inda/home).

## Codes Availability

1. Code used for GSMM Analysis of sulfur-starved *M. tuberculosis*: https://github.com/samrat-lab/Sulfur_Metabolism_Mtb

## Statistics and Reproducibility

The data values are represented as mean ± SEM of the replicates from two to three independent experiments. Wherever applicable, the number of replicates per group has been indicated by the dots. Statistical comparisons have been made with *Rv-WT (NT)* in most cases, unless otherwise noted. GraphPad Prism (ver 9.0) has been used for the statistical analysis of all data, with appropriate statistical tests and exact p-values reported in the respective figure legends.

## Acknowledgements

The authors acknowledge BRIC-THSTI for infrastructural support, the Experimental Animal Facility (and staff) for support with animal experiments, and the BSL-3 facility (and staff) for all *M. tuberculosis*-related experiments. Technical support of Mr. Sharad Dwivedi is duly acknowledged. We acknowledge Dr Prabhakar Babele, Multi-Omics facility, THSTI, for proteomics, and Dr B. N. Panda and Prabhanjan Dwivedi, Histopathology Facility, THSTI, for histopathological analysis of infected lung tissues. We acknowledge Indian Biological Data Centre (IBDC) for RNA-Seq Analysis. We duly acknowledge the support of Dr Ranjan K. Nanda, ICGEB, New Delhi, for ICPMS-based experiments. We also acknowledge receipt of the plasmids and constructs used in the study (Supplementary Information). This study was supported by the grant (No. EM/Dev/SG/212/7864/2023) from the Indian Council of Medical Research (ICMR), Government of India and intramural funding by BRIC-THSTI to AKP and is duly acknowledged. VKN was supported by a research fellowship from ICMR (3/1/3/JRF-2018/HRD-043/64467). VB is supported by the research fellowship from CSIR: 09/1049(11494)/2021-EMR-I. AM was supported by the research fellowship from ICMR: (3/1/3/JRF-2023/HRD-68/141819).

## Authors’ Contribution

VKN conceived of the study; VKN and AKP designed the research; VKN, VB, and AM performed BSL-3, microbiology-related experiments; MP and RP assisted with animal experiments in ABSL-3. GA and SC performed Genome Scale Metabolic Modelling (GSMM); VKN compiled the data; VKN, SC, RN and AKP analysed the data; VKN, GA, SC and AKP drafted; VKN and AKP revised the manuscript; AKP: funding acquisition and overall supervision. All authors reviewed and approved the final version.

## Additional Information

1. Supplementary Figures: Supplementary Figures and Legends

2. Supplementary Information: List of strains, plasmids, primers used in the study and other information.

## References

1. W. H. Organization, Bacterial Priority Pathogens List (WHO BPPL) (World Health Organization, 2024).

2. World Health Organization., “Global Tuberculosis Report 2025.” (2025).

3. C. Nathan, Mycobacterium tuberculosis as teacher. Nat Microbiol 8, 1606–1608 (2023).

4. A. Gouzy, Y. Poquet, O. Neyrolles, Nitrogen metabolism in Mycobacterium tuberculosis physiology and virulence. Nat Rev Microbiol 12, 729–737 (2014).

5. A. Gouzy, G. Larrouy-Maumus, D. Bottai, F. Levillain, A. Dumas, J. B. Wallach, I. Caire-Brandli, C. de Chastellier, T. D. Wu, R. Poincloux, R. Brosch, J. L. Guerquin-Kern, D. Schnappinger, L. P. Sorio de Carvalho, Y. Poquet, O. Neyrolles, Mycobacterium tuberculosis exploits asparagine to assimilate nitrogen and resist acid stress during infection. PLoS Pathog 10, e1003928 (2014).

6. A. Gouzy, G. Larrouy-Maumus, T. D. Wu, A. Peixoto, F. Levillain, G. Lugo-Villarino, J. L. Guerquin-Kern, L. P. de Carvalho, Y. Poquet, O. Neyrolles, Mycobacterium tuberculosis nitrogen assimilation and host colonization require aspartate. Nat Chem Biol 9, 674–676 (2013).

7. A. Agapova, A. Serafini, M. Petridis, D. M. Hunt, A. Garza-Garcia, C. D. Sohaskey, L. P. S. de Carvalho, Flexible nitrogen utilisation by the metabolic generalist pathogen Mycobacterium tuberculosis. Elife 8 (2019).

8. K. Borah, M. Beyss, A. Theorell, H. Wu, P. Basu, T. A. Mendum, K. Nӧh, D. J. V Beste, J. McFadden, Intracellular Mycobacterium tuberculosis Exploits Multiple Host Nitrogen Sources during Growth in Human Macrophages. Cell Rep 29, 3580–3591 e4 (2019).

9. R. M. Gray, D. M. Hunt, M. Silva Dos Santos, J. Liu, A. Agapova, A. Rodgers, A. Fearns, J. O. Canseco, A. Garza-Garcia, J. I. MacRae, M. G. Gutierrez, R. E. Lee, L. P. S. de Carvalho, Mycobacterium tuberculosis overcomes phosphate starvation by extensively remodelling its lipidome with phosphorus-free lipids. Nat Commun 16, 11317 (2025).

10. C. Healy, S. Ehrt, A. Gouzy, An exacerbated phosphate starvation response triggers Mycobacterium tuberculosis glycerol utilization at acidic pH. mBio 16, e0282524 (2025).

11. S. Ehrt, D. Schnappinger, K. Y. Rhee, Metabolic principles of persistence and pathogenicity in Mycobacterium tuberculosis. Nat Rev Microbiol 16, 496–507 (2018).

12. L. Huang, E. V Nazarova, D. G. Russell, Mycobacterium tuberculosis: Bacterial Fitness within the Host Macrophage. Microbiol Spectr 7 (2019).

13. G. T. Mashabela, T. J. de Wet, D. F. Warner, Mycobacterium tuberculosis Metabolism. Microbiol Spectr 7 (2019).

14. Michael W. Schelle and Carolyn R Bertozzi, Sulfate Metabolism in Mycobacteria. ChemBioChem 7, 1516–1524 (2006).

15. V. Saini, B. M. Cumming, L. Guidry, D. A. Lamprecht, J. H. Adamson, V. P. Reddy, K. C. Chinta, J. H. Mazorodze, J. N. Glasgow, M. Richard-Greenblatt, A. Gomez-Velasco, H. Bach, Y. Av-Gay, H. Eoh, K. Rhee, A. J. C. Steyn, Ergothioneine Maintains Redox and Bioenergetic Homeostasis Essential for Drug Susceptibility and Virulence of Mycobacterium tuberculosis. Cell Rep 14, 572–585 (2016).

16. M. A. Rahman, B. M. Cumming, K. W. Addicott, H. T. Pacl, S. L. Russell, K. Nargan, T. Naidoo, P. K. Ramdial, J. H. Adamson, R. Wang, A. J. C. Steyn, Hydrogen sulfide dysregulates the immune response by suppressing central carbon metabolism to promote tuberculosis. Proc Natl Acad Sci U S A 117, 6663–6674 (2020).

17. V. Saini, K. C. Chinta, V. P. Reddy, J. N. Glasgow, A. Stein, D. A. Lamprecht, M. A. Rahman, J. S. Mackenzie, B. E. Truebody, J. H. Adamson, T. T. R. Kunota, S. M. Bailey, D. R. Moellering, J. R. Lancaster Jr., A. J. C. Steyn, Hydrogen sulfide stimulates Mycobacterium tuberculosis respiration, growth and pathogenesis. Nat Commun 11, 557 (2020).

18. M. D. Howe, S. L. Kordus, M. S. Cole, A. A. Bauman, C. C. Aldrich, A. D. Baughn, Y. Minato, Methionine Antagonizes para-Aminosalicylic Acid Activity via Affecting Folate Precursor Biosynthesis in Mycobacterium tuberculosis. Front Cell Infect Microbiol 8, 399 (2018).

19. P. J. Kies, N. D. Hammer, A Resourceful Race: Bacterial Scavenging of Host Sulfur Metabolism during Colonization. Infect Immun 90, e0057921 (2022).

20. C. Lai, L. Yang, V. Pathiranage, R. Wang, F. V Subach, A. R. Walker, K. D. Piatkevich, Genetically encoded green fluorescent sensor for probing sulfate transport activity of solute carrier family 26 member a2 (Slc26a2) protein. Commun Biol 7, 1375 (2024).

21. W. P. Gill, N. S. Harik, M. R. Whiddon, R. P. Liao, J. E. Mittler, D. R. Sherman, A replication clock for Mycobacterium tuberculosis. Nat Med 15, 211–214 (2009).

22. S. Helaine, J. A. Thompson, K. G. Watson, M. Liu, C. Boyle, D. W. Holden, Dynamics of intracellular bacterial replication at the single cell level. Proc Natl Acad Sci U S A 107, 3746–3751 (2010).

23. J. M. Mouton, S. Helaine, D. W. Holden, S. L. Sampson, Elucidating population-wide mycobacterial replication dynamics at the single-cell level. Microbiology (Reading) 162, 966–978 (2016).

24. E. S. Kavvas, Y. Seif, J. T. Yurkovich, C. Norsigian, S. Poudel, W. W. Greenwald, S. Ghatak, B. O. Palsson, J. M. Monk, Updated and standardized genome-scale reconstruction of Mycobacterium tuberculosis H37Rv, iEK1011, simulates flux states indicative of physiological conditions. BMC Syst Biol 12, 25 (2018).

25. B. Ofori-Anyinam, M. Hamblin, M. L. Coldren, B. Li, G. Mereddy, M. Shaikh, A. Shah, C. Grady, N. Ranu, S. Lu, P. C. Blainey, S. Ma, J. J. Collins, J. H. Yang, Catalase activity deficiency sensitizes multidrug-resistant Mycobacterium tuberculosis to the ATP synthase inhibitor bedaquiline. Nat Commun 15, 9792 (2024).

26. C. Colijn, A. Brandes, J. Zucker, D. S. Lun, B. Weiner, M. R. Farhat, T. Y. Cheng, D. B. Moody, M. Murray, J. E. Galagan, Interpreting expression data with metabolic flux models: predicting Mycobacterium tuberculosis mycolic acid production. PLoS Comput Biol 5, e1000489 (2009).

27. J. Schellenberger, B. O. Palsson, Use of randomized sampling for analysis of metabolic networks. J Biol Chem 284, 5457–5461 (2009).

28. H. A. Herrmann, B. C. Dyson, L. Vass, G. N. Johnson, J. M. Schwartz, Flux sampling is a powerful tool to study metabolism under changing environmental conditions. NPJ Syst Biol Appl 5, 32 (2019).

29. S. A. Bhat, I. K. Iqbal, A. Kumar, Imaging the NADH:NAD(+) Homeostasis for Understanding the Metabolic Response of Mycobacterium to Physiologically Relevant Stresses. Front Cell Infect Microbiol 6, 145 (2016).

30. T. Sharma, S. Tyagi, R. Pal, J. Kundu, S. K. Gupta, V. Barik, V. K. Nain, M. Pandey, P. Dwivedi, B. N. Panda, Y. Kumar, R. K. Nanda, S. Chatterjee, A. K. Pandey, Phosphoglucomutase A-mediated metabolic adaptation is essential for antibiotic and disease persistence in *Mycobacterium tuberculosis*. mSystems 10, 1–23 (2025).

31. E. Wooff, S. L. Michell, S. V Gordon, M. A. Chambers, S. Bardarov, W. R. Jacobs Jr., R. G. Hewinson, P. R. Wheeler, Functional genomics reveals the sole sulphate transporter of the Mycobacterium tuberculosis complex and its relevance to the acquisition of sulphur in vivo. Mol Microbiol 43, 653–663 (2002).

32. A. S. Zolotarev, M. Unnikrishnan, B. E. Shmukler, J. S. Clark, D. H. Vandorpe, N. Grigorieff, E. J. Rubin, S. L. Alper, Increased sulfate uptake by E. coli overexpressing the SLC26-related SulP protein Rv1739c from Mycobacterium tuberculosis. Comp Biochem Physiol A Mol Integr Physiol 149, 255–266 (2008).

33. J. M. Rock, F. F. Hopkins, A. Chavez, M. Diallo, M. R. Chase, E. R. Gerrick, J. R. Pritchard, G. M. Church, E. J. Rubin, C. M. Sassetti, D. Schnappinger, S. M. Fortune, Programmable transcriptional repression in mycobacteria using an orthogonal CRISPR interference platform. Nat Microbiol 2, 16274 (2017).

34. A. I. Wong, J. M. Rock, CRISPR Interference (CRISPRi) for Targeted Gene Silencing in Mycobacteria. Methods Mol Biol 2314, 343–364 (2021).

35. E. Aguilar-Barajas, C. Diaz-Perez, M. I. Ramirez-Diaz, H. Riveros-Rosas, C. Cervantes, Bacterial transport of sulfate, molybdate, and related oxyanions. Biometals 24, 687–707 (2011).

36. S. Ehrt, Controlling gene expression in mycobacteria with anhydrotetracycline and Tet repressor. Nucleic Acids Res. 33, e21–e21 (2005).

37. M. A. Rahman, J. N. Glasgow, S. Nadeem, V. P. Reddy, R. R. Sevalkar, J. R. Lancaster Jr., A. J. C. Steyn, The Role of Host-Generated H2S in Microbial Pathogenesis: New Perspectives on Tuberculosis. Front Cell Infect Microbiol 10, 586923 (2020).

38. V. M. Brunner, P. W. Fowler, Compensatory mutations are associated with increased in vitro growth in resistant clinical samples of Mycobacterium tuberculosis. Microb. Genom. 10 (2024).

39. W. Worakitchanon, H. Yanai, P. Piboonsiri, R. Miyahara, S. Nedsuwan, W. Imsanguan, B. Chaiyasirinroje, W. Sawaengdee, S. Wattanapokayakit, N. Wichukchinda, Y. Omae, P. Palittapongarnpim, K. Tokunaga, S. Mahasirimongkol, A. Fujimoto, Comprehensive analysis of Mycobacterium tuberculosis genomes reveals genetic variations in bacterial virulence. Cell Host Microbe 32, 1972–1987.e6 (2024).

40. The CRyPTIC Consortium, A data compendium associating the genomes of 12,289 Mycobacterium tuberculosis isolates with quantitative resistance phenotypes to 13 antibiotics. PLoS Biol. 20, e3001721 (2022).

41. P. The CRyPTIC Consortium & Fowler, The CRyPTIC Consortium Dataset (Version v3.4.0). [Preprint] (2025). 10.5281/zenodo.15680920.

42. Broad Institute (BI), SRX470553: Illumina whole genome shotgun sequencing of genomic DNA paired-end library “Solexa-215485” containing sample “XTB13-198,” https://www.ncbi.nlm.nih.gov/sra/SRX470553[accn] (2014).

43. W. Le Mouellic, F. Levillain, T. D. Wu, M. Caouaille, P. Bousso, Y. Poquet, O. Neyrolles, Inorganic sulfate is critical for Mycobacterium tuberculosis lung tissue colonization and redox balance. Proc Natl Acad Sci U S A 122, e2503966122 (2025).

44. J. C. Betts, P. T. Lukey, L. C. Robb, R. A. McAdam, K. Duncan, Evaluation of a nutrient starvation model of Mycobacterium tuberculosis persistence by gene and protein expression profiling. Mol Microbiol 43, 717–731 (2002).

45. M. Gengenbacher, S. P. S. Rao, K. Pethe, T. Dick, Nutrient-starved, non-replicating Mycobacterium tuberculosis requires respiration, ATP synthase and isocitrate lyase for maintenance of ATP homeostasis and viability. Microbiology (Reading) 156, 81– 87 (2010).

46. E. A. Weinstein, T. Yano, L. S. Li, D. Avarbock, A. Avarbock, D. Helm, A. A. McColm, K. Duncan, J. T. Lonsdale, H. Rubin, Inhibitors of type II NADH:menaquinone oxidoreductase represent a class of antitubercular drugs. Proc Natl Acad Sci U S A 102, 4548–4553 (2005).

47. T. Beites, K. O’Brien, D. Tiwari, C. A. Engelhart, S. Walters, J. Andrews, H. J. Yang, M. L. Sutphen, D. M. Weiner, E. K. Dayao, M. Zimmerman, B. Prideaux, P. V Desai, T. Masquelin, L. E. Via, V. Dartois, H. I. Boshoff, C. E. Barry 3rd, S. Ehrt, D. Schnappinger, Plasticity of the Mycobacterium tuberculosis respiratory chain and its impact on tuberculosis drug development. Nat Commun 10, 4970 (2019).

48. C. Vilcheze, B. Weinrick, L. W. Leung, W. R. Jacobs Jr., Plasticity of Mycobacterium tuberculosis NADH dehydrogenases and their role in virulence. Proc Natl Acad Sci U S A 115, 1599–1604 (2018).

49. A. Pathak, S. Saini, R. Pathania, Disruption of cysteine metabolism leads to synthetic lethality and in vivo fitness impairment in Acinetobacter baumannii. mBio, e0084226 (2026).

50. T. Tralau, S. Vuilleumier, C. Thibault, B. J. Campbell, C. A. Hart, M. A. Kertesz, Transcriptomic analysis of the sulfate starvation response of Pseudomonas aeruginosa. J Bacteriol 189, 6743–6750 (2007).

51. J. E. Dominy Jr., C. R. Simmons, P. A. Karplus, A. M. Gehring, M. H. Stipanuk, Identification and characterization of bacterial cysteine dioxygenases: a new route of cysteine degradation for eubacteria. J Bacteriol 188, 5561–5569 (2006).

52. E. P. Tchesnokov, M. Fellner, E. Siakkou, T. Kleffmann, L. W. Martin, S. Aloi, I. L. Lamont, S. M. Wilbanks, G. N. Jameson, The cysteine dioxygenase homologue from Pseudomonas aeruginosa is a 3-mercaptopropionate dioxygenase. J Biol Chem 290, 24424–24437 (2015).

53. K. Shatalin, E. Shatalina, A. Mironov, E. Nudler, H2S: a universal defense against antibiotics in bacteria. Science (1979). 334, 986–990 (2011).

54. C. Vilcheze, T. R. Weisbrod, B. Chen, L. Kremer, M. H. Hazbon, F. Wang, D. Alland, J. C. Sacchettini, W. R. Jacobs Jr., Altered NADH/NAD+ ratio mediates coresistance to isoniazid and ethionamide in mycobacteria. Antimicrob Agents Chemother 49, 708–720 (2005).

55. M. Nguyen, A. Quemard, S. Broussy, J. Bernadou, B. Meunier, Mn(III) pyrophosphate as an efficient tool for studying the mode of action of isoniazid on the InhA protein of Mycobacterium tuberculosis. Antimicrob Agents Chemother 46, 2137–2144 (2002).

56. I. Levin-Reisman, I. Ronin, O. Gefen, I. Braniss, N. Shoresh, N. Q. Balaban, Antibiotic tolerance facilitates the evolution of resistance. Science (1979). 355, 826–830 (2017).

57. X. Wang, W. J. Jowsey, C. Y. Cheung, N. Dickerhof, C. L. Champan, J. R. H. Taka, M. B. Hampton, G. Bashiri, P. P. Gardner, P. C. Fineran, G. M. Cook, S. A. Jackson, M. B. McNeil, Genome scale CRISPRi reveals both shared and strain-specific vulnerabilities in genetically diverse drug-resistant strains of Mycobacterium tuberculosis. Nat Commun, doi: 10.1038/s41467-026-73952-x (2026).

58. V. K. Nain, “Sulfur metabolism and its role in mycobacterial physiology and pathogenesis,” thesis, BRIC-Translational Health Science and Technology Institute, Faridabad and Jawarharlal Nehru University, New Delhi, India, New Delhi (2024).

59. P. R. Wheeler, N. G. Coldham, L. Keating, S. V Gordon, E. E. Wooff, T. Parish, R. G. Hewinson, Functional demonstration of reverse transsulfuration in the Mycobacterium tuberculosis complex reveals that methionine is the preferred sulfur source for pathogenic Mycobacteria. J Biol Chem 280, 8069–8078 (2005).

60. G. H. Babunovic, M. A. DeJesus, B. Bosch, M. R. Chase, T. Barbier, A. K. Dickey, B. D. Bryson, J. M. Rock, S. M. Fortune, CRISPR Interference Reveals That All-Trans-Retinoic Acid Promotes Macrophage Control of Mycobacterium tuberculosis by Limiting Bacterial Access to Cholesterol and Propionyl Coenzyme A. mBio, e0368321 (2022).

61. B. Bosch, M. A. DeJesus, N. C. Poulton, W. Zhang, C. A. Engelhart, A. Zaveri, S. Lavalette, N. Ruecker, C. Trujillo, J. B. Wallach, S. Li, S. Ehrt, B. T. Chait, D. Schnappinger, J. M. Rock, Genome-wide gene expression tuning reveals diverse vulnerabilities of M. tuberculosis. Cell 184, 4579–4592 e24 (2021).

62. T. D. Schmittgen, K. J. Livak, Analyzing real-time PCR data by the comparative C(T) method. Nat Protoc 3, 1101–1108 (2008).

63. R. Pal, S. Talwar, M. Pandey, V. K. Nain, T. Sharma, S. Tyagi, V. Barik, S. Chaudhary, S. K. Gupta, Y. Kumar, R. Nanda, A. Singhal, A. K. Pandey, Rv0495c regulates redox homeostasis in Mycobacterium tuberculosis. Tuberculosis 145, 102477–102489 (2024).

64. M. I. Love, W. Huber, S. Anders, Moderated estimation of fold change and dispersion for RNA-seq data with DESeq2. Genome Biol 15, 550 (2014).

65. G. Arora, M. Banerjee, J. Langthasa, R. Bhat, S. Chatterjee, Targeting metabolic fluxes reverts metastatic transitions in ovarian cancer. iScience 26, 108081 (2023).

66. L. Heirendt, S. Arreckx, T. Pfau, S. N. Mendoza, A. Richelle, A. Heinken, H. S. Haraldsdottir, J. Wachowiak, S. M. Keating, V. Vlasov, S. Magnusdottir, C. Y. Ng, G. Preciat, A. Zagare, S. H. J. Chan, M. K. Aurich, C. M. Clancy, J. Modamio, J. T. Sauls, A. Noronha, A. Bordbar, B. Cousins, D. C. El Assal, L. V Valcarcel, I. Apaolaza, S. Ghaderi, M. Ahookhosh, M. Ben Guebila, A. Kostromins, N. Sompairac, H. M. Le, D. Ma, Y. Sun, L. Wang, J. T. Yurkovich, M. A. P. Oliveira, P. T. Vuong, L. P. El Assal, I. Kuperstein, A. Zinovyev, H. S. Hinton, W. A. Bryant, F. J. Aragon Artacho, F. J. Planes, E. Stalidzans, A. Maass, S. Vempala, M. Hucka, M. A. Saunders, C. D. Maranas, N. E. Lewis, T. Sauter, B. O. Palsson, I. Thiele, R. M. T. Fleming, Creation and analysis of biochemical constraint-based models using the COBRA Toolbox v.3.0. Nat Protoc 14, 639–702 (2019).

67. A. Gupta, A. Kumar, R. Anand, N. Bairagi, S. Chatterjee, Genome scale metabolic model driven strategy to delineate host response to Mycobacterium tuberculosis infection. Mol Omics 17, 296–306 (2021).

68. G. Bachrach, M. J. Colston, H. Bercovier, D. Bar-Nir, C. Anderson, K. G. Papavinasasundaram, A new single-copy mycobacterial plasmid, pMF1, from Mycobacterium fortuitum which is compatible with the pAL5000 replicon. Microbiology (Reading) 146 **(Pt 2)**, 297–303 (2000).

69. Y. Cai, E. Jaecklein, J. S. Mackenzie, K. Papavinasasundaram, A. J. Olive, X. Chen, A. J. C. Steyn, C. M. Sassetti, Host immunity increases Mycobacterium tuberculosis reliance on cytochrome bd oxidase. PLoS Pathog 17, e1008911 (2021).

70. S. Helaine, A. M. Cheverton, K. G. Watson, L. M. Faure, S. A. Matthews, D. W. Holden, Internalization of Salmonella by macrophages induces formation of nonreplicating persisters. Science 343, 204–8 (2014).

71. C. Michaux, S. Ronneau, S. Helaine, Studying Antibiotic Persistence During Infection. Methods Mol Biol 2357, 273–289 (2021).

72. S. Ronneau, C. Michaux, S. Helaine, Decline in nitrosative stress drives antibiotic persister regrowth during infection. Cell Host Microbe 31, 993–1006 e6 (2023).

73. P. Babele, R. B. Kumar, S. Rajoria, F. Rashid, D. Malakar, S. S. Bhagyawant, D. V Kamboj, S. I. Alam, Putative serum protein biomarkers for epsilon toxin exposure in mouse model using LC-MS/MS analysis. Anaerobe 63, 102209 (2020).

74. C. Chen, Y. Wu, J. Li, X. Wang, Z. Zeng, J. Xu, Y. Liu, J. Feng, H. Chen, Y. He, R. Xia, TBtools-II: A “one for all, all for one” bioinformatics platform for biological big-data mining. Mol Plant 16, 1733–1742 (2023).

75. V. K. Nain, V. Barik, M. Pandey, M. Pareek, T. Sharma, R. Pal, S. Tyagi, M. Bajpai, P. Dwivedi, B. N. Panda, Y. Kumar, S. Asthana, A. K. Pandey, A pH-dependent direct sulfhydrylation pathway is required for the pathogenesis of Mycobacterium tuberculosis. *Commun*. Biol. 8, 637–652 (2025).

