## Supplementary Figures and Information for "Orthogonal perturbation of sulfur availability reveals antibiotic-induced synthetic lethality in *Mycobacterium tuberculosis*"

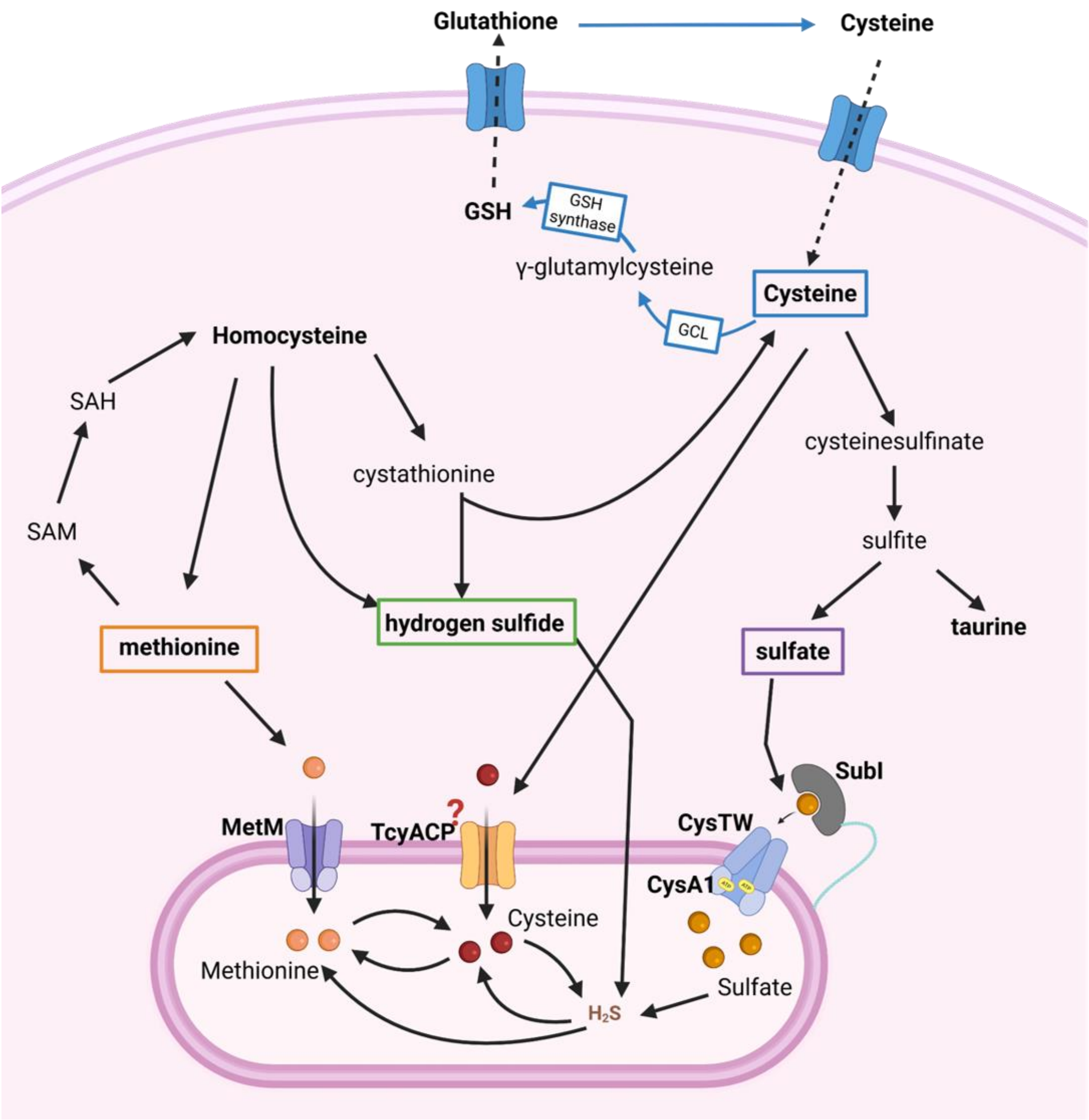


**Fig. S1: Acquisition of inorganic sulfate by *M. tuberculosis* and sulfur dependent biological processes.** *M. tuberculosis* uptakes the primary S-source in the form of inorganic sulfate (Si) via ABC transporter CysTWA1-SubI or in the form of organic sources like cysteine and methionine through dedicated transporters. Bacteria can also use hydrogen sulfide ($H_2S$) as sulfur source to re-fuel or recharge its thiol pool and does not require any dedicated transporter. All these nutrients are converted into one form or the other to sustain survival. While on other hand, the host acquire sulfur in the form of reduced sources like cysteine, methionine, glutathione, etc.

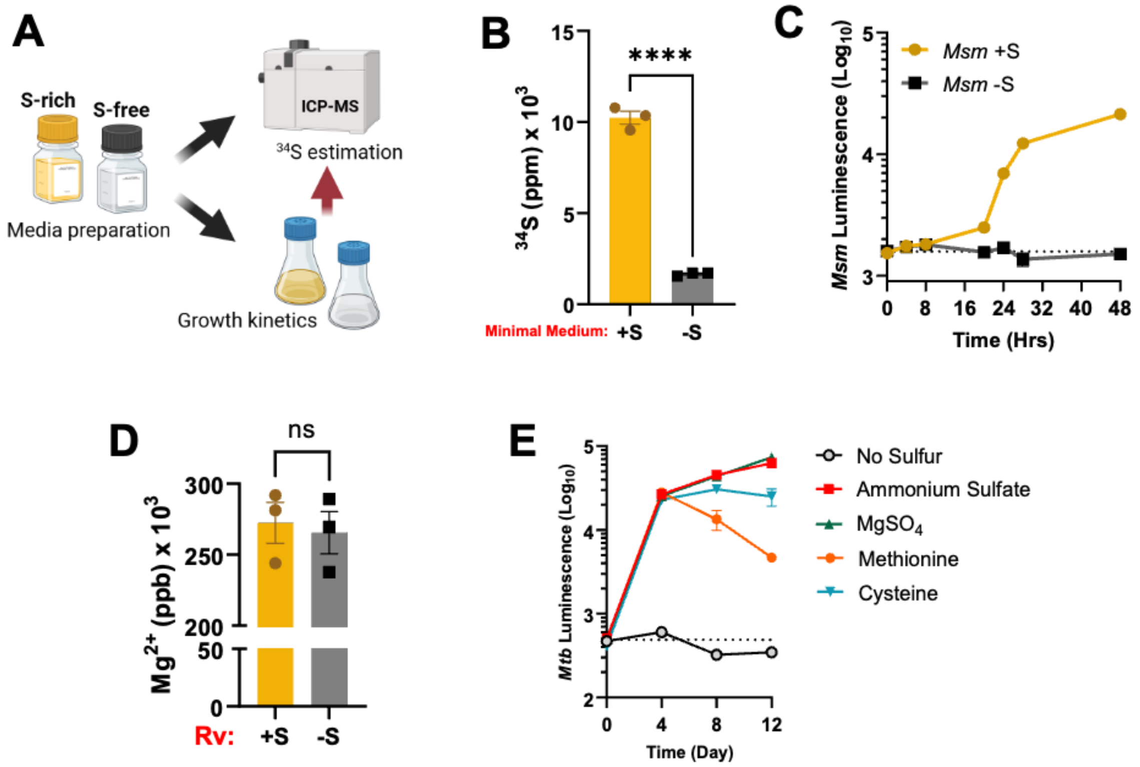


**Fig. S2: Exclusion of inorganic sulfur hampers mycobacterial growth. (A)** Schematic representation showing preparation of chemically defined sulfur-rich and sulfur-free media, verified by ICP-MS for total sulfur content, followed by growth kinetics of mycobacteria in the respective media. **(B)** ICP-MS was used to estimate $^{34}S$ inorganic sulfur in chemically defined sulfur-rich and sulfur-free media. **(C)** Growth kinetics of *M. smegmatis* in sulfur-rich and sulfur-free media. **(D)** ICP-MS for assay of Mg levels in *M. tuberculosis* grown in minimal medium with or without sulfur. **(E)** *M. tuberculosis* cultured in sulfur-free medium and supplemented with various sources of sulfur, inorganic as well as amino acids. Data are the averages of the three data points indicated by dots and representative of at least two to three independent experiments. Error bars correspond to standard error of the mean (SEM). Statistical significance was assessed by Unpaired Student's t-test for (B) and (D), where ns, non-significant and ****P<0.0001.

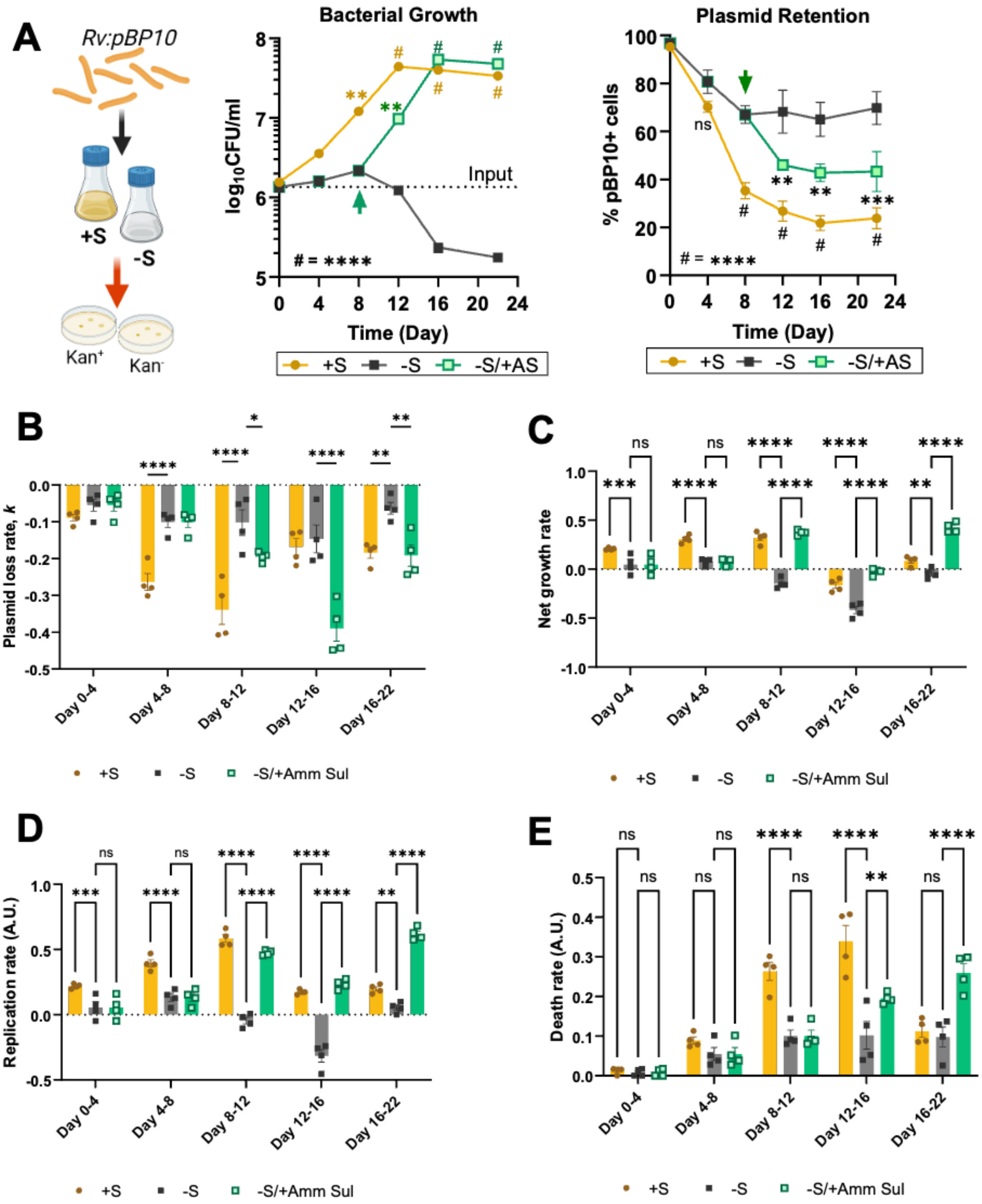


**Fig. S3: Replication kinetics of *M. tuberculosis* grown under S-depleted conditions. (A)** Determining the replication rate of *M. tuberculosis* in the presence and absence of sulfur using the replication clock plasmid, pBP10 indicating growth and plasmid retention at indicated time points. **(B)** Plasmid loss rate, *k* **(C)** Net growth rate, *n* **(D)** Replication rate, *r* **(E)** Death rate, *d* as calculated for the respective bacterial cultures for the time periods as indicated. Data are the averages of the three data points indicated by dots and representative of at least two to three independent experiments. Error bars correspond to standard error of mean (SEM). Statistical significance was assessed by two-way ANOVA (Dunnett's multiple comparisons test) for (A-E), where, ns, non-significant; *P<0.05; **P<0.01; ***P<0.001 and ****P<0.0001S

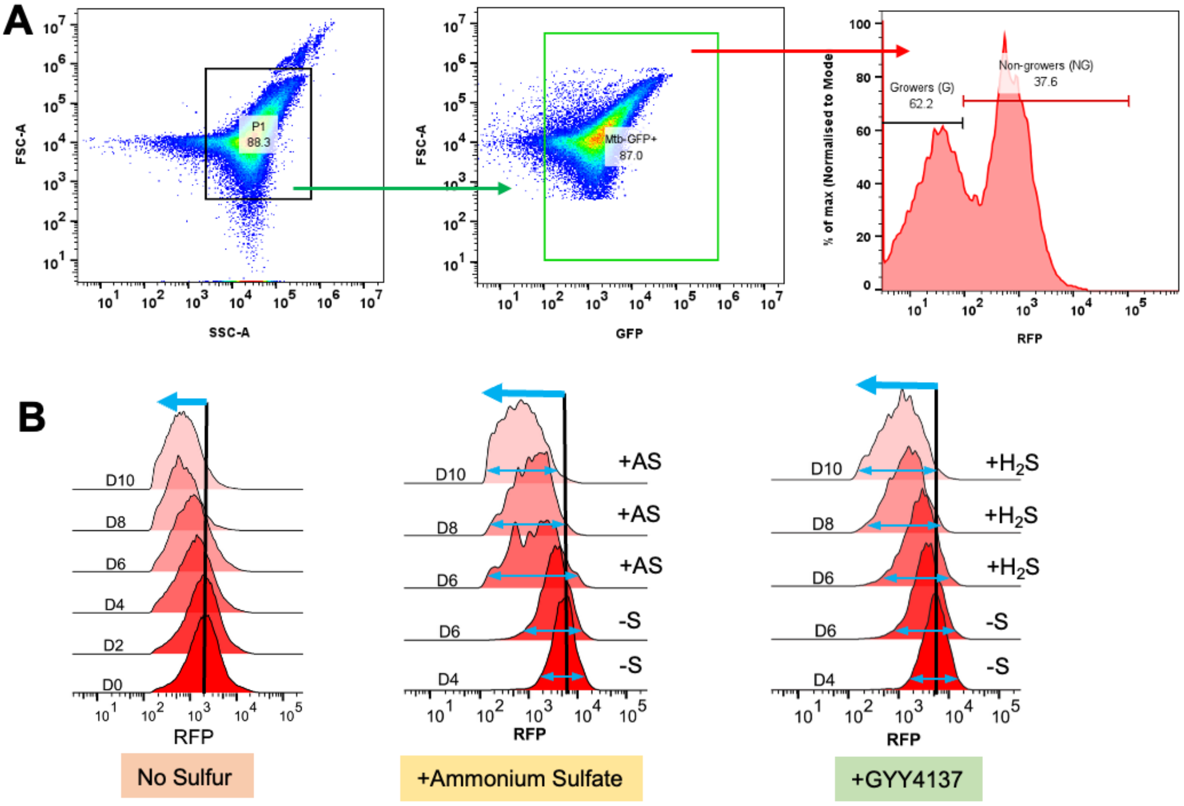


**Fig. S4: Fluorescence dilution reveals enrichment of non-growers upon S-limitation in *M. tuberculosis*. (A)** Gating strategy for analysing GFP+ cells with variable RFP expression. GFP was used to distinguish cells from debris, while RFP served as a replication marker. **(B)** Histograms depicting dilution of RFP with time and upon supplementation of sulfur sources – Ammonium sulfate (inorganic) or GYY4137 ($H_2S$-donor, gaseous).

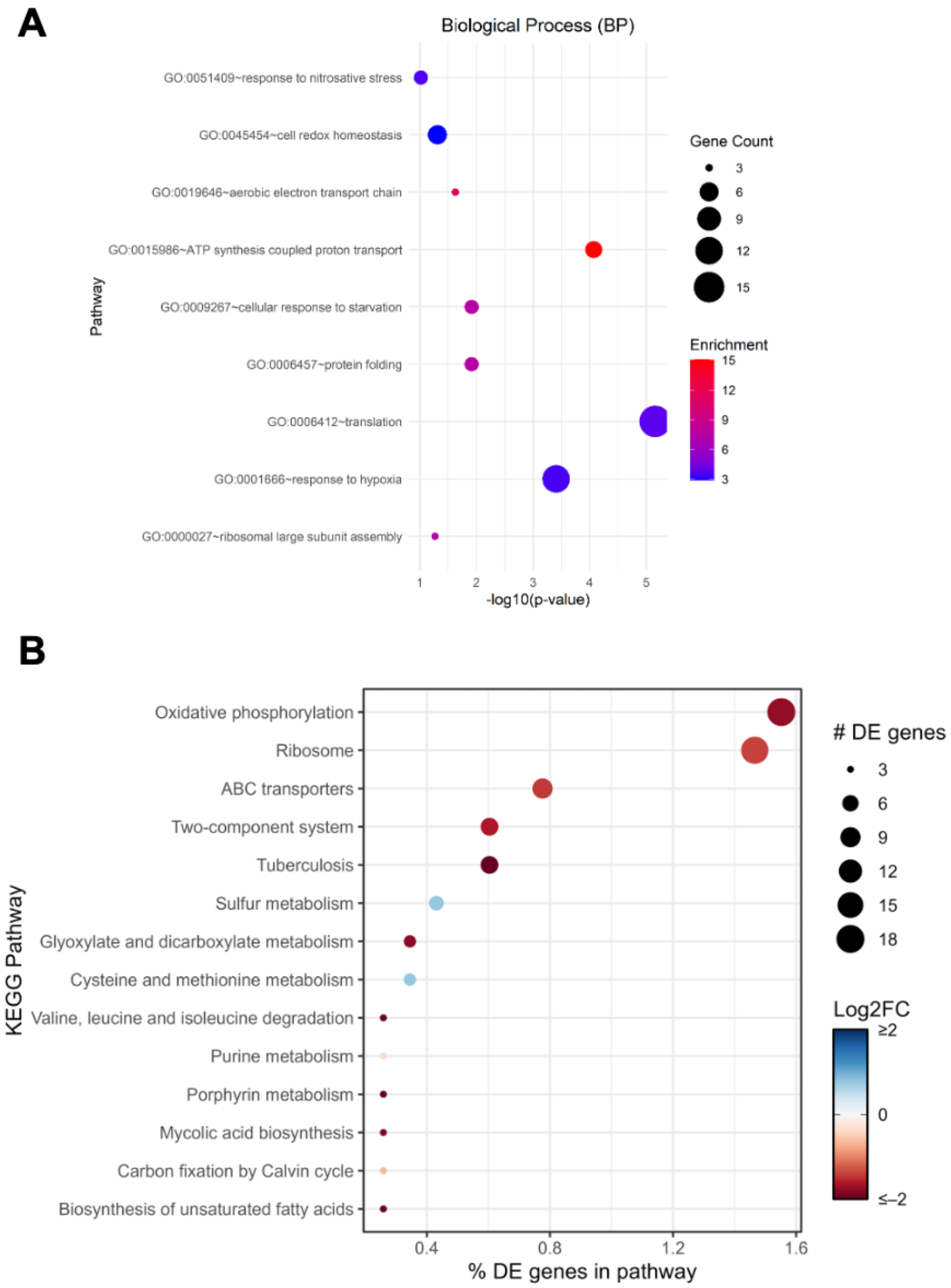


**Fig. S5: Gene Ontology terms and pathway analysis of S-starved *M. tuberculosis*. (A)** Gene Ontology (GO) terms associated with biological processes (BP) affected by S-starvation. **(B)** KEGG Pathway analysis for metabolic and physiological pathways affected by significant expression of genes involved in respective processes.

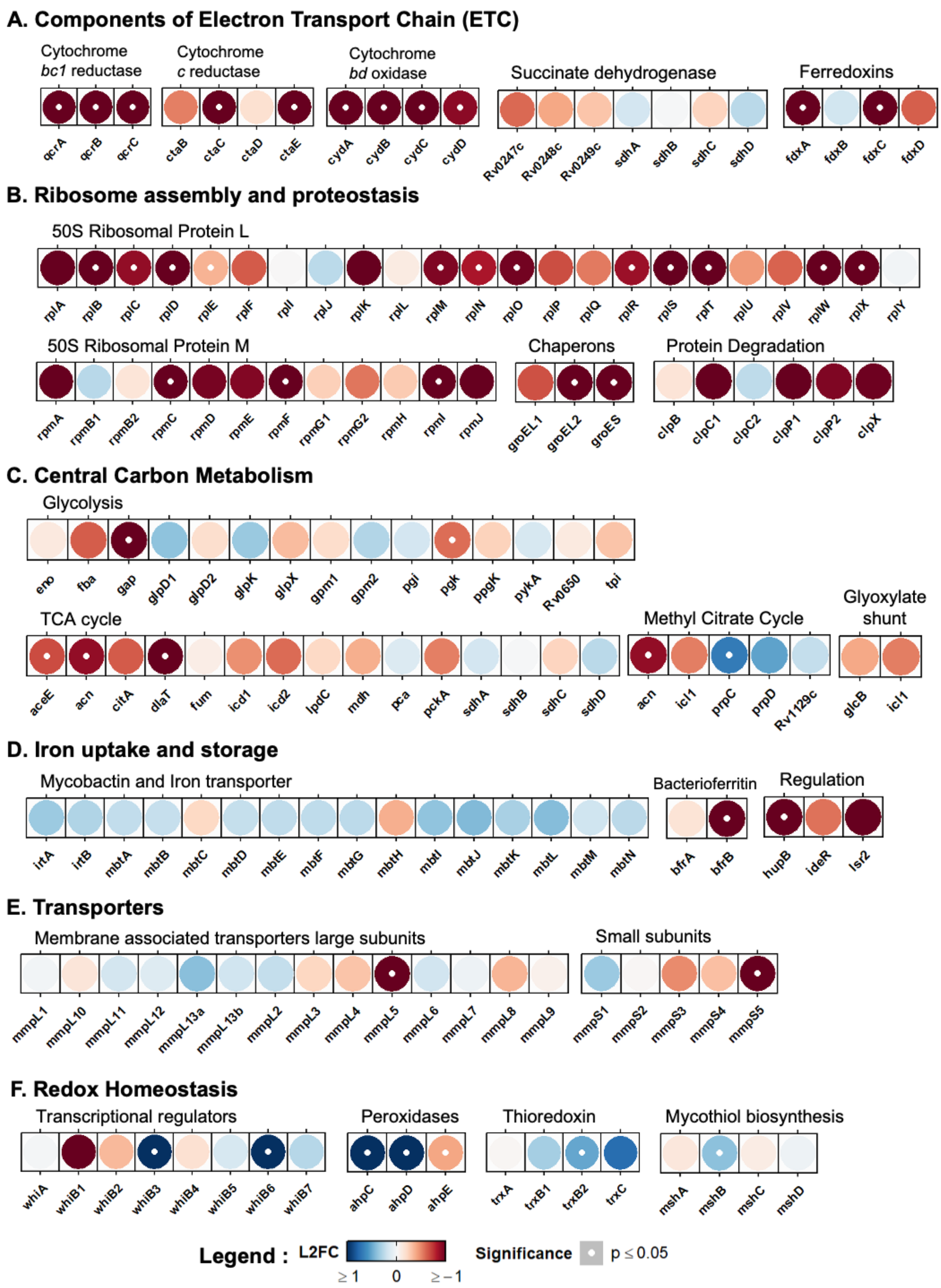
A. Components of Electron Transport Chain (ETC)
Cytochrome bc1 reductase
Cytochrome c reductase
Cytochrome bd oxidase
Succinate dehydrogenase
Ferredoxins
qcrA qcrB qcrC ctaB ctaC ctaD ctaE cydA cydB cydC cydD Rv0247c Rv0248c Rv0249c sdhA sdhB sdhC sdhD fdxA fdxB fdxC fdxD
B. Ribosome assembly and proteostasis
50S Ribosomal Protein L
rplA rplB rplC rplD rplE rplF rplI rplJ rplK rplL rplM rplN rplO rplP rplQ rplR rplS rplT rplU rplV rplW rplX rplY
50S Ribosomal Protein M
Chaperons
Protein Degradation
rpmA rpmB1 rpmB2 rpmC rpmD rpmE rpmF rpmG1 rpmG2 rpmH rpmI rpmJ groEL1 groEL2 groES clpB clpC1 clpC2 clpP1 clpP2 clpX
C. Central Carbon Metabolism
Glycolysis
eno fba gap glpD1 glpD2 glpK glpX gpm1 gpm2 pgi pgk ppgK pykA Rv0650 tpi
TCA cycle
Methyl Citrate Cycle
Glyoxylate shunt
aceE acn citA dlaT fum icd1 icd2 lpdC mdh pca pckA sdhA sdhB sdhC sdhD acn icl1 prpC prpD Rv1129c glcB icl1
D. Iron uptake and storage
Mycobactin and Iron transporter
Bacterioferritin
Regulation
irtA irtB mbtA mbtB mbtC mbtD mbtE mbtF mbtG mbtH mbtI mbtJ mbtK mbtL mbtM mbtN bfrA bfrB hupB ideR lsr2
E. Transporters
Membrane associated transporters large subunits
Small subunits
mmpL1 mmpL10 mmpL11 mmpL12 mmpL13a mmpL13b mmpL2 mmpL3 mmpL4 mmpL5 mmpL6 mmpL7 mmpL8 mmpL9 mmpS1 mmpS2 mmpS3 mmpS4 mmpS5
F. Redox Homeostasis
Transcriptional regulators
Peroxidases
Thioredoxin
Mycothiol biosynthesis
whiA whiB1 whiB2 whiB3 whiB4 whiB5 whiB6 whiB7 ahpC ahpD ahpE trxA trxB1 trxB2 trxC mshA mshB mshC mshD
Legend : L2FC
≥ 1 0 ≥ −1
Significance
p ≤ 0.05

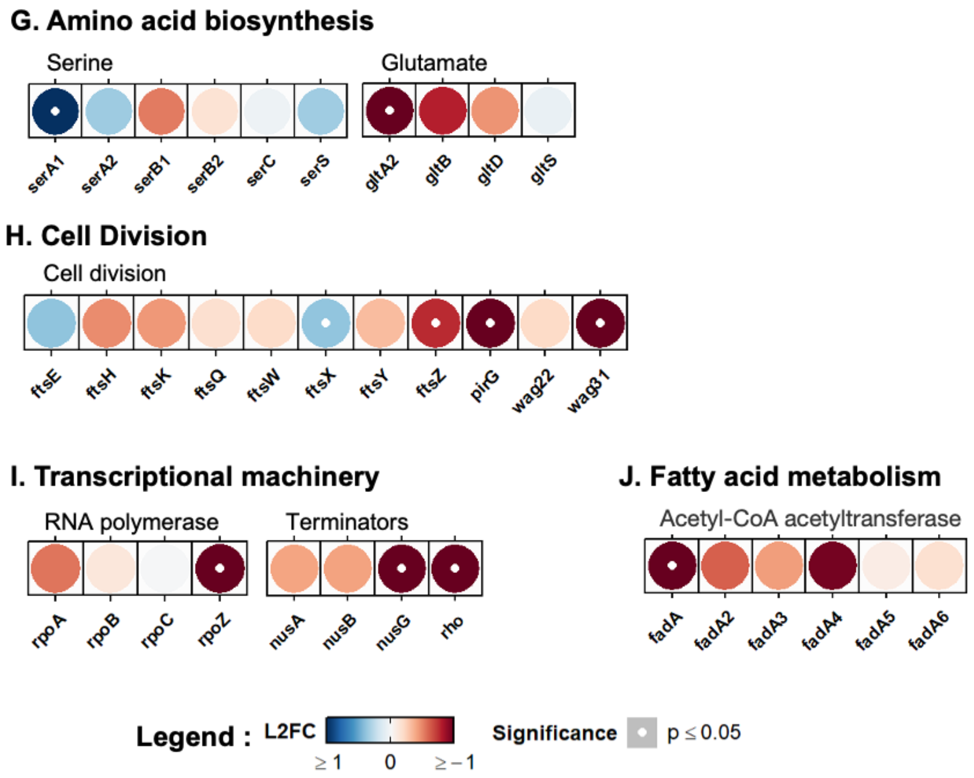


**Fig. S6: Feature expression heatmap of the DEGs involved in different physiological pathways in response to sulfur depletion in *M. tuberculosis*.** The color of each circle represents the expression levels in $\log_2$ fold change (L2FC); a white dot represents p-value (Significance) ≤ 0.05 with |L2FC|>1 as cut-off for change in gene expression denoted as upregulation or downregulated. **(A)** Components of Electron transport chain (ETC) –Cytochrome Q reductase, Cytochrome *bd* oxidase, succinate dehydrogenases and ferredoxins; **(B)** Ribosome assembly and protein homeostasis–50S (*rpl* and *rpm*) ribosome subunit proteins, chaperonins and protein degradation machinery; **(C)** Central carbon metabolism – glycolysis, TCA cycle, methyl citrate cycle (MCC) and Glyoxylate shunt; **(D)** Iron acquisition (*mbt*), uptake (*irt*), storage (*bfr*) and regulation (transcriptional regulators); **(E)** Membrane transporters – membrane-associated transporters (*mmpLS*); **(F)** Redox homeostasis – redox-dependent transcription factors (*whiB* family), alkyl hydroperoxidases, thioredoxin reductases, mycothiol biosynthesis etc.; **(G)** Amino acid biosynthesis – serine, and glutamate; **(H)** Cell division associated genes; **(I)** Transcriptional machinery – genes encoding subunits of RNA Polymerase, transcriptional terminators – (Nus and Rho) and **(J)** Fatty acid metabolism – a family of *fad* genes involved in acetyl transfer.

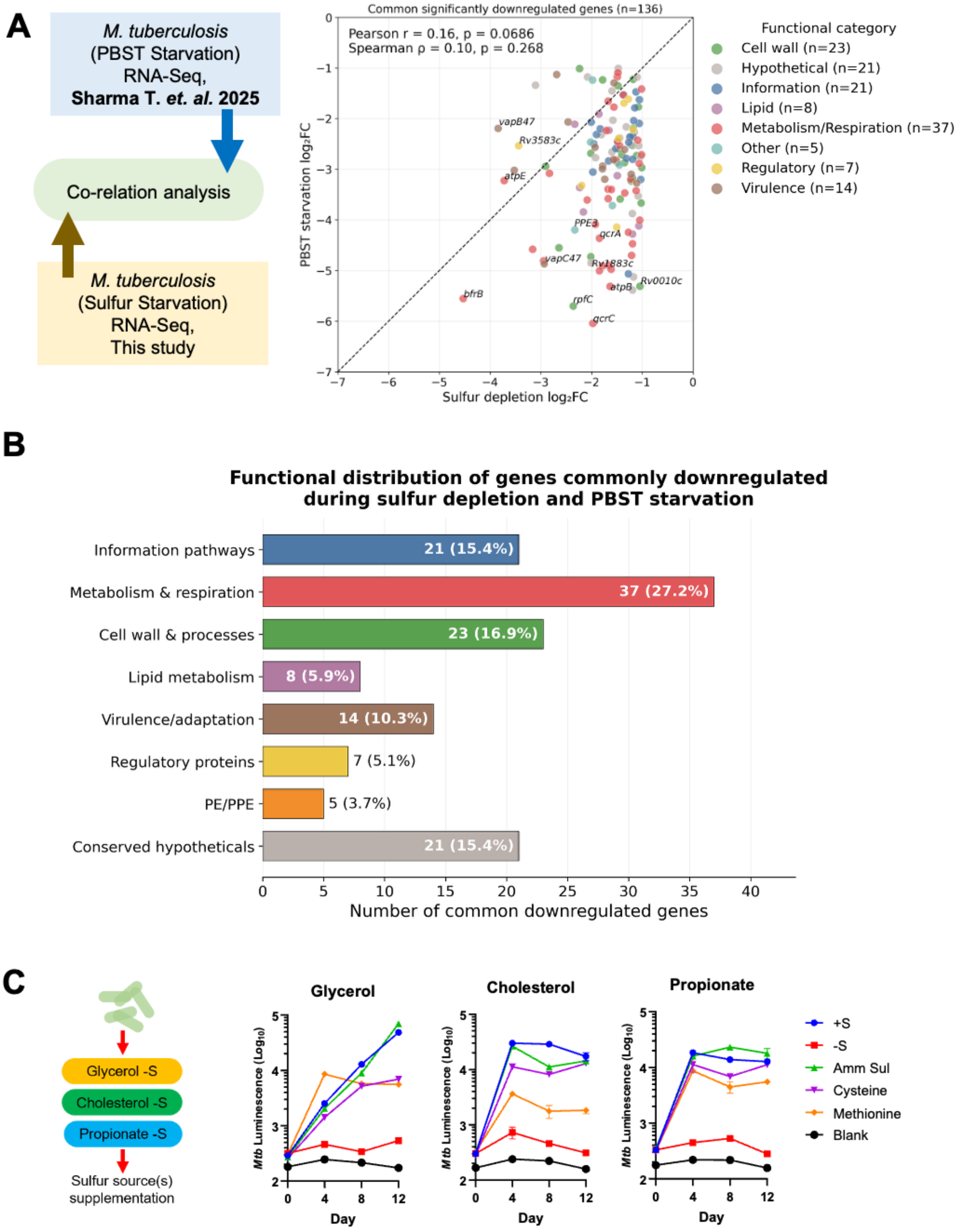


**Fig. S7: Comparison of the transcriptomic profile of sulfur and nutrient-starved *M. tuberculosis*. (A)** Transcriptomic profiles of sulfur-starved and nutrient-starved (in PBST) *M. tuberculosis* were compared to identify common up- and downregulated genes. Expression (L2FC) across both conditions was correlated, and significantly up- and down-regulated genes (L2FC ≥ 1 and *p*-adjusted value ≤ 0.05) were highlighted. **(B)** Functional categorisation of genes commonly down-regulated in both conditions. **(C)** Growth kinetics of *M. tuberculosis* in the presence and absence of inorganic sulfur with different carbon sources – 0.2% glycerol, 200μM cholesterol and 10mM propionate. S-starved bacteria were supplemented with different sources of sulfur – inorganic (ammonium sulfate) and organic (cysteine, methionine).

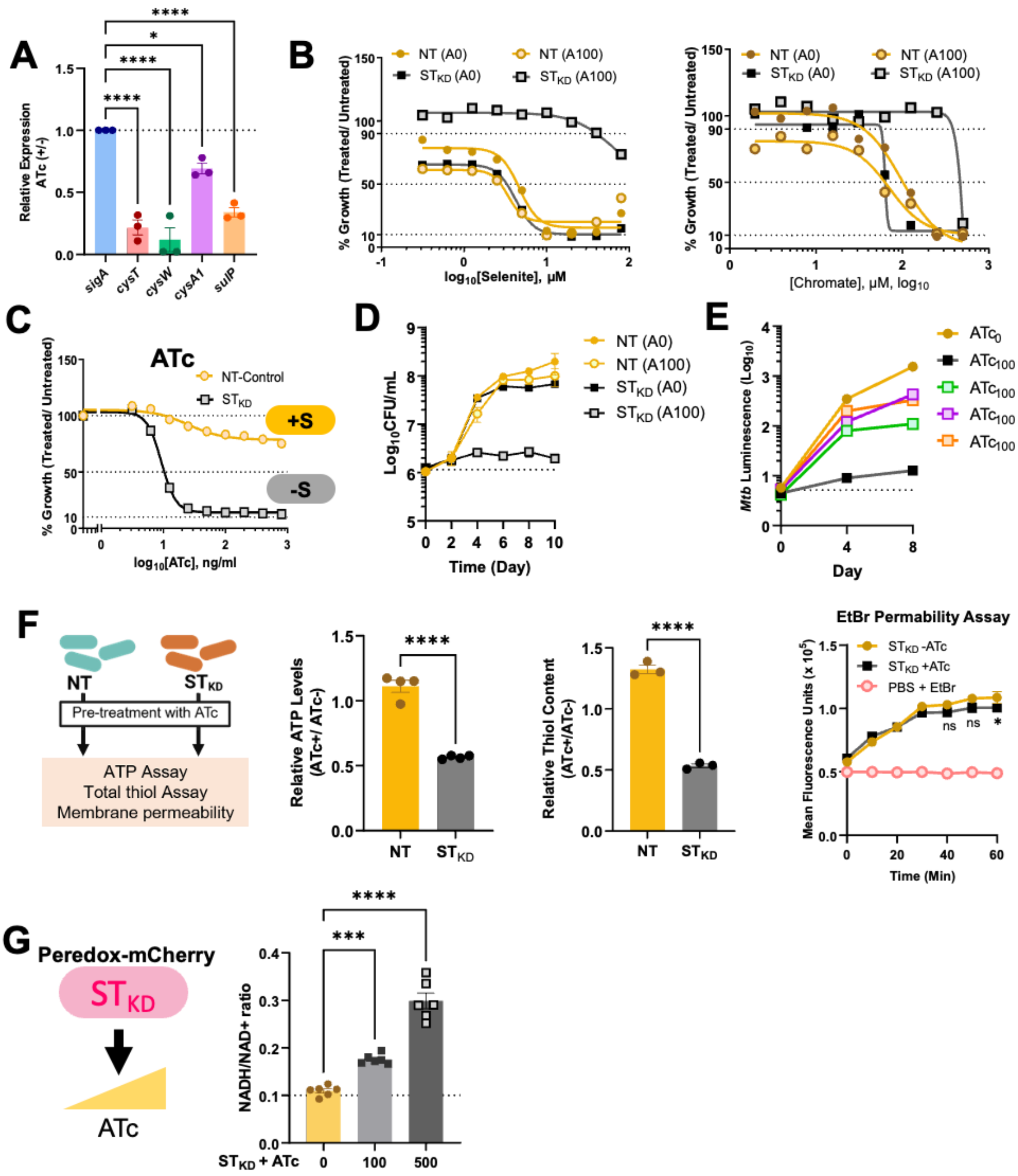


**Fig. S8: Transcriptional knockdown of sulfate transporter (ST) encoding genes and its metabolic consequences. (A)** Relative expression of target genes encoding sulfate transporter(s) upon knockdown of CRISPRi by Anhydrotetracycline (ATc) confirmed by RT-qPCR with *sigA* as internal control **(B)** Dose-response for selenium and chromate resistance. **(C)** ATc titration assay for defining vulnerability in *M. tuberculosis* in response to knockdown of sulfate transporter(s). **(D)** Growth kinetics of NT and $ST_{KD}$ *M. tuberculosis*. **(E)** Growth kinetics of *M. tuberculosis* lacking sulfate transporter ($ST_{KD}$) supplemented with different sulfur sources – NaHS, cysteine and methionine. **(F)** $ST_{KD}$ (transporter knockdown) and NT (control) strains were treated with ATc to induce CRISPRi mediated knockdown followed by respective biochemical assays – ATP, total thiol and membrane permeability. **(G)** $ST_{KD}$ *M. tuberculosis* expressing Peredox-mCherry biosensor to assess intracellular NADH/NAD+ ratio upon knockdown of sulfate transporter(s). Data are the averages of the three data points indicated by dots and representative of at least two to three independent experiments. Error bars correspond to standard error of mean (SEM). Statistical significance was assessed by Ordinary one-way ANOVA (Dunnett's multiple comparisons test) for (A), where, *P=0.0110 and ****P<0.0001; Unpaired Student's t-test for (F-ATP and thiol assay), where, ****P<0.0001; two-way ANOVA (Sidak's multiple comparisons test) for (F-EtBr assay), where, *P = 0.0356 and Ordinary one-way ANOVA (Dunnett's multiple comparisons test) for (G), where, ***P = 0.0006 and ****P<0.0001.

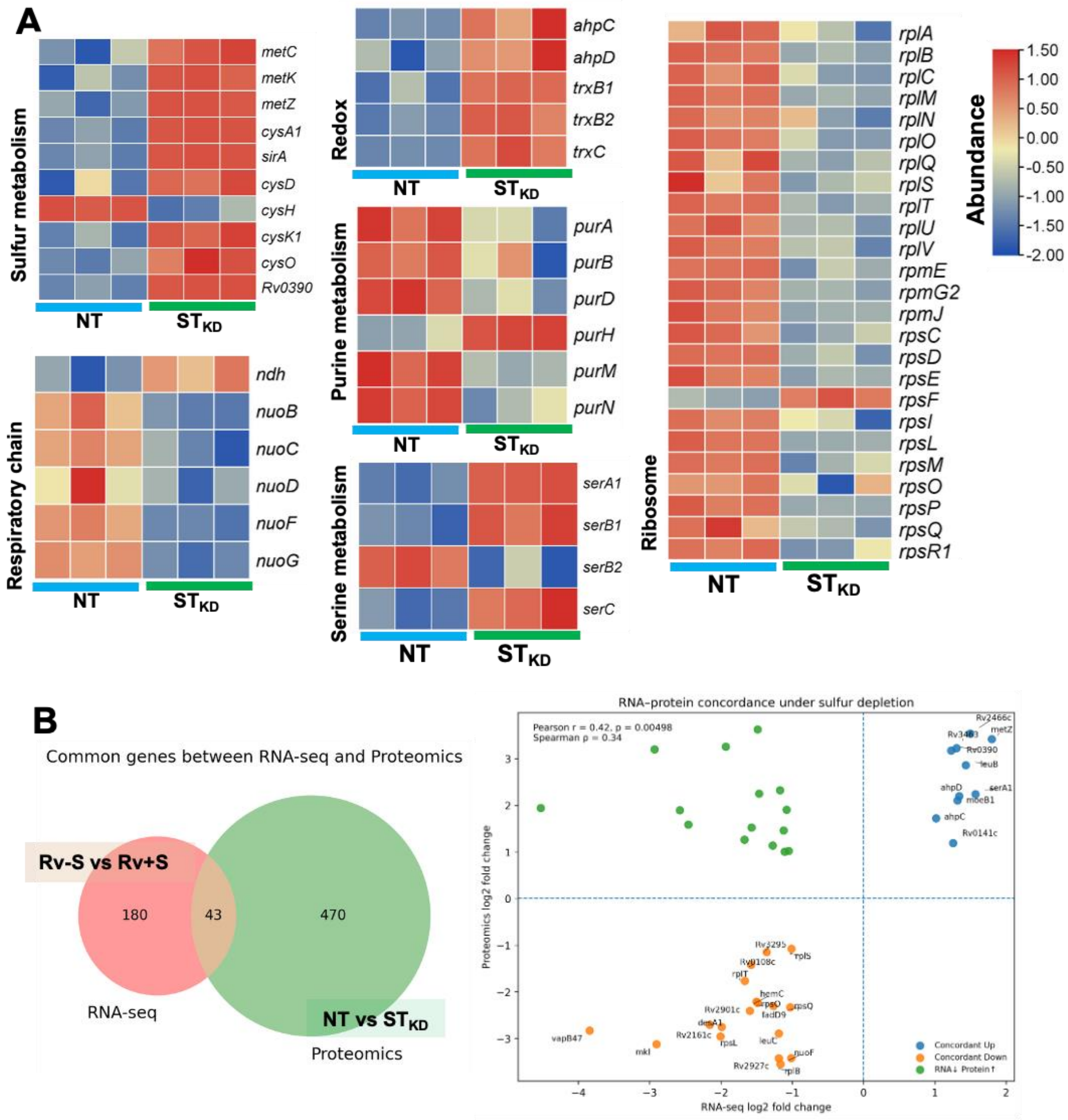


**Fig. S9: Proteomic analysis of sulfate transporter-lacking *M. tuberculosis*. (A)** Heatmaps showing representative functional categories with altered protein abundance in $ST_{KD}$. **(B)** Correlation of S-starved *M. tuberculosis* transcriptomics and $ST_{KD}$ *M. tuberculosis* proteomics for analysis of concordant transcripts and proteins.

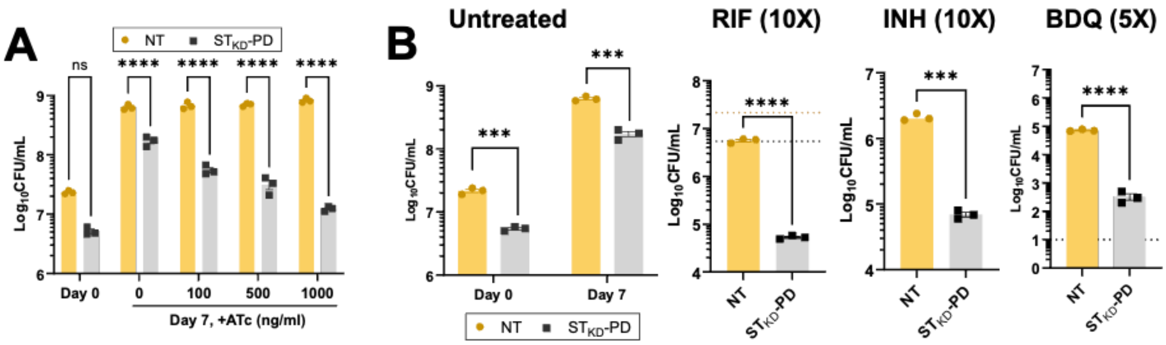


**Fig. S10: In-vitro drug efficacy assay.** NT and $ST_{KD}$ *M. tuberculosis* were pre-treated with ATc to deplete sulfate transporter transcripts and treated with **(A)** ATc (0, 100, 500, and 1000 ng/ml) and **(B)** $ATc_{100}$ in combination with anti-TB drugs at higher concentrations for 7 days. Bacterial viability was determined by CFU plating. The data are the averages of the three data points indicated by dots and are representative of at least three independent experiments. Error bars correspond to standard error of mean (SEM). Statistical significance was assessed by two-way ANOVA (Sidak's multiple comparisons test) for (A), where ****P<0.0001 and Unpaired Student's t-test for (B), where ***P = 0.0002 and ****P<0.0001.

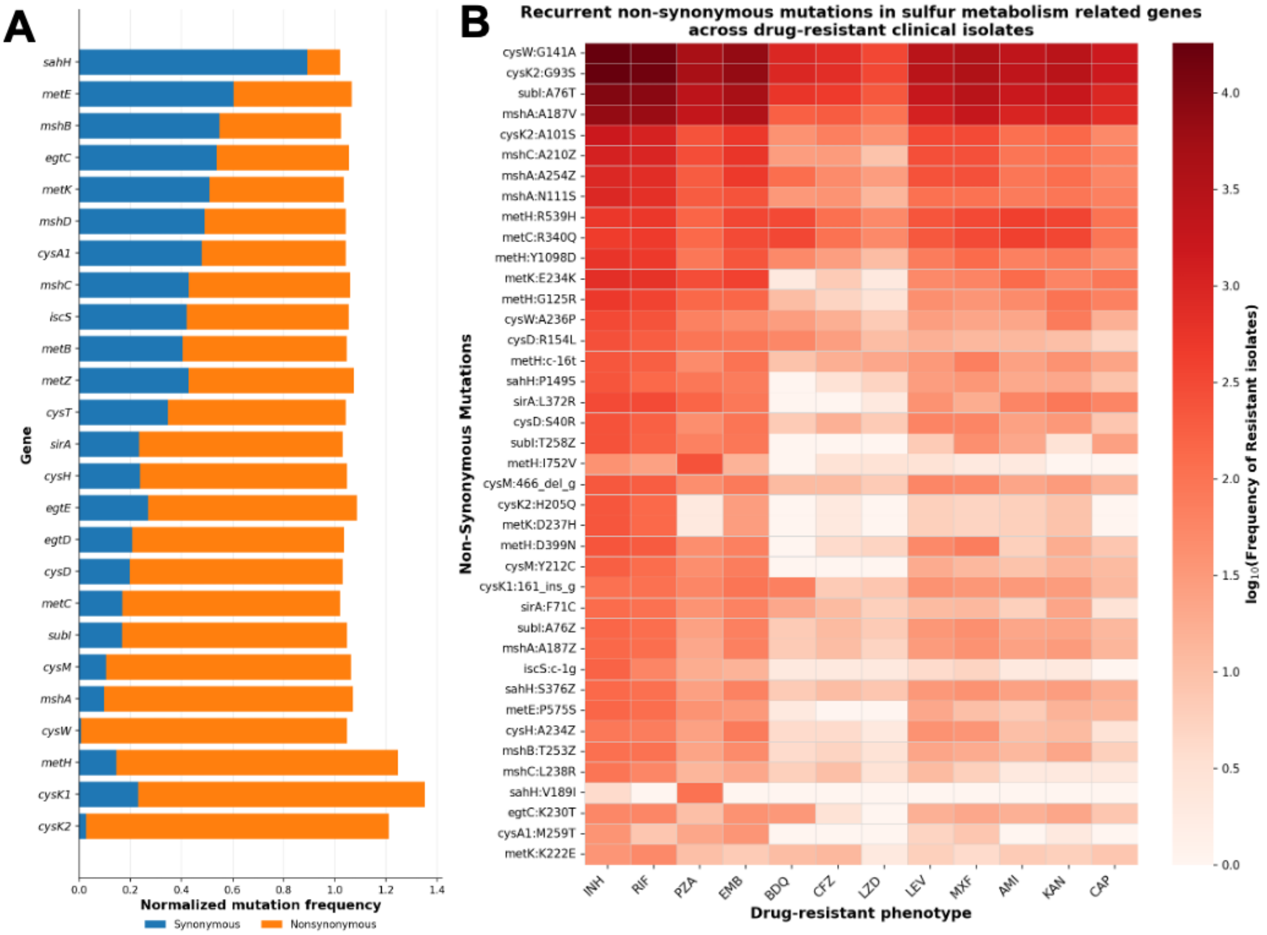


**Fig. S11: Genomic analysis of clinical isolates of *M. tuberculosis* from the CRyPTIC database. (A)** Analysis depicting normalised synonymous vs non-synonymous mutations in the genes involved in sulfur metabolism. **(B)** Heatmap showing the frequently occurring non-synonymous mutations in sulfur metabolism-related genes.

**List of plasmid constructs used in the study**

| # | Plasmid | Feature | Reference |
|---|---|---|---|
| 1. | pIJR965 | Kan, *attB* integrative, target sgRNA and *dCas9*$_{Sth}$ expressing plasmid for transcriptional knockdown of target gene(s) in *M. tuberculosis* | Kind gift from Prof. Sarah M. Fortune |
| 2. | pIJR965::ctrlsgRNA | Control guide expressing CRISPRi construct | This study |
| 3. | pIJR965::*cysT* guide | *cysT* targeting CRISPRi construct | This study |
| 4. | pIJR965::*cysW* guide | *cysW* targeting CRISPRi construct | This study |
| 5. | pIJR965::*rv1739c* guide | *rv1739c (sulP)* targeting CRISPRi construct | This study |
| 6 | pIJR965::*cysT_cysW_rv1739c* guides | Tandemly *cysT, cysW* and *rv1739c* targeting CRISPRi construct | This study |
| 7. | pMV261: mEmerald, TagRFP | Hyg, Mycobacterial Live-Dead reporter with ATc inducible RFP | Kind gift from Prof. Christopher M. Sassetti |
| 8. | pJW276 | Zeo, Integrative, Lux expressing plasmid | Kind gift from Prof. Sarah M. Fortune |
| 9. | pIMT100 (pMV762:Peredox-mCherry) | Hyg, NADH/NAD+ biosensor expressing plasmid | Kind gift from Dr. Ashwani Kumar, CSIR-IMTECH, India |
| 10. | pST-H | Hyg, Episomal mycobacterial expression vector | Kind gift from Dr. Vinay K. Nandicoori, CSIR-CCMB, India |
| 11. | pST-H:Thyone-tagRFP (mThyone) | Hyg, Sulfate biosensor expressing plasmid | This study |
| 12. | pBP10 | Kan, Mycobacterial replication clock plasmid | Gill *et. al.* 2009 |

**List of strains used in the study**

| S. No. | Strain | Marker | Feature | Reference |
|---|---|---|---|---|
| 1. | *E. coli* XL1Blue | | | This lab |
| 2. | *M. tuberculosis* $H_{37}Rv$ (*Rv*) | | Wild type *Mtb (Rv-WT)* | Kind gift from Christopher M. Sassetti |
| 3. | *Rv*::*lux* | *Zeo*$^r$ | *Rv*-WT Luciferase reporter | Nain VK *et. al.* 2025 |
| 4. | *Rv::lux:pBP10* | *Zeo*$^r$ *Kan*$^r$ | *Rv*-WT Luciferase reporter harboring replication clock plasmid | This work |
| 5. | *Rv*: pMV261 mEmerald, TagRFP | *Hyg*$^r$ | Rv-WT expressing dual reporter (GFP+RFP_ | Nain VK *et. al.* 2025 |
| 6. | *Rv*:pIMT100 (Peredox-mCherry) | *Hyg*$^r$ | Rv-WT expressing NADH/NAD+ ratiometric biosensor | This work |
| 7. | *Rv*:pST-H *mThyone* | *Hyg*$^r$ | Rv-WT expressing sulfate ratiometric biosensor | This work |
| 8. | *Rv::lux::NT* | *Zeo*$^r$ *Kan*$^r$ | Rv-WT with control sgRNA (non-targeting) CRISPRi control luminescence reporter strain | Nain VK *et. al.* 2025 |
| 9. | *Rv::lux::cysTW+sulP*$_{KD}$ ($ST_{KD}$) | *Zeo*$^r$ *Kan*$^r$ | Rv-WT with *cysT* + *cysW* and *sulP* targeting CRISPRi luminescence reporter strain | This work |
| 10. | $ST_{KD}$:pIMT100 (Peredox-mCherry) | *Zeo*$^r$ *Kan*$^r$ *Hyg*$^r$ | $ST_{KD}$: strain expressing NADH/NAD+ biosensor | This work |
| 11. | $ST_{KD}$:pST-H *mThyone* | *Zeo*$^r$ *Kan*$^r$ *Hyg*$^r$ | $ST_{KD}$: strain expressing sulfate biosensor | This work |
| 12. | XTB13-198 ($INH^R$) | $INH^r$ | Isoniazid resistant clinical *M. tuberculosis* isolate | BEI Resources, NIAID, NIH (CAT: NR-49364) |
| 13. | XTB13-198 ($INH^R$) :: *lux* | $INH^r$ $Zeo^r$ | Isoniazid resistant clinical *M. tuberculosis* isolate luminescence reporter strain | This work |
| 14. | XTB13-198 ($INH^R$) :: *lux:: ST*$_{KD}$ | $INH^r$ $Zeo^r$ $Kan^r$ | Isoniazid resistant clinical *M. tuberculosis* isolate luminescence reporter strain with sulfate transporters targeted using CRISPRi | This work |

**List of Primers used in the study**

| # | Primer Name | Sequence 5'-3' | Feature |
|---|---|---|---|
| 1. | *cysT*-CR-F | GGGAGCGGCATGCGACGAGACCGCCAG | CRISPRi guide for *cysT* |
| 2. | *cysT*-CR-R | AAACCTGGCGGTCTCGTCGCATGCCGC | |
| 3. | *cysW*-CR-F | GGGAACCGGATGGAGGGCAGCGTGATTCG | CRISPRi guide for *cysW* |
| 4. | *cysW*-CR-R | AAACCGAATCACGCTGCCCTCCATCCGGT | |
| 5. | *Rv1739c*-CR-F | GGGAGCGAAAGAGTGTACTTCGCT | CRISPRi guide for *Rv1739c* |
| 6. | *Rv1739c*-CR-R | AAACAGCGAAGTACACTCTTTCGC | |
| 7. | *cysT*-RT-F | GGTCATCAACCTGGTGTTCGG | qPCR primers of *cysT* |
| 8. | *cysT*-RT-R | GAACGGCAATGTGACGAACGC | |
| 9. | *cysW*-RT-F | CACGACGGCATTGGTGCTG | qPCR primers of *cysW* |
| 10. | *cysW*-RT-R | ACCACGAATGGACAGGTGACG | |
| 11. | *subI*-RT-F | CCGACCTGGTGAACTTCTCG | qPCR primers of *subI* |
| 12. | *subI*-RT-R | GATTCCACTTGGCAGAACCCG | |
| 13. | *cysA1*-RT-F | CATCGGATTCGTCTTCCAGCAC | qPCR primers of *cysA1* |
| 14. | *cysA1*-RT-R | CGAGCAGCAGCACCTCC | |
| 15. | *sigA*-RT-F | GAAGACCACGAAGACCTCGAAG | qPCR primers of *sigA* |
| 16. | *sigA*-RT-R | GTTTGAGGTAGGCGCGAACC | |
| 17. | *Rv1739c*-RT-F | AGCCAACTCGGCACTATCACC | qPCR primers of *rv1739c* |
| 18. | *Rv1739c*-RT-R | CCGACCCACAATCGCAATACCTTT | |
| 19. | GG_Vector-F | TATGCTCTTCAGGATCTGACCAGGGAAAA TAGCC | Primers for amplifying sgRNA cassettes |
| 20. | GG-Vector-R | TATGCTCTTCACTGAAAATAAAAAAGGGG ACCTCTAGGG | |
| 21. | GG-Vector-F2 | TATGCTCTTCAGATAAAATAAAAAAGGGG ACCTCTAGGG | |
| 22. | GG-Vector-R1 | TATGCTCTTCAATCTCTGACCAGGGAAAA TAGCC | |
